# Nischarin agonist rilmenidine shows antimetastatic potential in pancreatic ductal adenocarcinoma

**DOI:** 10.1101/2025.10.27.684770

**Authors:** Kristina Živić, Marijana Pavlović, Marija Ostojić, Danijel Galun, Aleksandar Pavić, Tatjana Srdić-Rajić, Jelena Grahovac

## Abstract

Pancreatic ductal adenocarcinoma (PDAC) is one of the most aggressive cancer types with a dismal prognosis. Early metastatic dissemination and rich desmoplastic stroma limit therapeutic efficacy, and the discovery of new treatment approaches that will target these obstacles is of great importance. Nischarin (NISCH)/Imidazoline-1 receptor (IR-1) is a potential tumor suppressor, that is involved in the regulation of cell migration, invasion, and cytoskeletal organization. Importantly, several FDA-approved agonists target this receptor. This study aimed to examine the expression of NISCH in PDAC and the effects of its agonists with the intent of drug repurposing. NISCH was expressed in PDAC tumor tissue, in both the epithelial and stromal compartments of tumors, and higher NISCH expression was associated with longer patient survival. Out of the three tested NISCH agonists, rilmenidine was the most potent and could induce cancer cell apoptosis. Rilmenidine treatment induced transcriptional changes in cancer cells associated with so far reported NISCH function and decreased their adhesion and migration *in vitro*. In CAFs, rilmenidine treatment decreased the expression of activation markers and limited cancer cell-CAF cytokine communication in co-culture. Ultimately, in the *in vivo* tumor xenograft zebrafish model, rilmenidine reduced the metastatic spread of pancreatic cancer cells. Our results suggest that NISCH agonist rilmenidine is a promising candidate for drug repurposing as an antimetastatic agent in PDAC and that NISCH can be a potential target for the development of new PDAC therapeutics.

## INTRODUCTION

Pancreatic cancer is one of the most lethal cancers with a 5-year survival rate of 11% (1,2). Pancreatic ductal adenocarcinoma (PDAC) accounts for almost 90% of all pancreatic cancer cases (3). With clinically silent early lesions, PDAC is generally diagnosed in an already advanced stage, when it has metastasized to other organs and the treatment options are limited (4). Only 20% of PDAC patients are amenable to potentially curable surgical removal (5). Unfortunately, even in operated patients the 5-year survival rate remains below 30% (6–9). Metastatic dissemination in PDAC starts very early (10) and it is estimated that only five years are required for the acquisition of the metastatic ability of the initial non-metastatic founder cell (11). Despite improvements in therapeutic approaches, PDAC remains poorly responsive to conventional, targeted, and immunotherapies (12–15), and currently there are no specific approaches for targeting the metastatic disease.

Fibrosis is the main feature of PDAC, and the dense stroma consisting of the extracellular matrix, sparse immune infiltrate, cancer-associated fibroblasts (CAFs), and tumor vasculature can comprise up to 90% of the tumor mass (16,17). CAFs with their heterogeneous nature participate in various signaling pathways that contribute to the tumor progression (18). Their role in the modification of the extracellular matrix (19), cancer cell migration and invasion processes (20), as well as the drug resistance (21–23), raised awareness about the necessity of examining the effects of systemically delivered drugs not only on cancer cells but also on the tumor microenvironment (TME).

Nischarin (NISCH) is a cytosolic protein, first identified in mice as a binding partner of integrin α5, involved in the regulation of cell motility (24). Soon after, it was recognized that it is homologous to the IRAS-I1 receptor antisera-selected protein in humans (25). Over the past 20 years it has been shown that NISCH has tumor suppressor properties in breast cancer, capable of modulating cell migration and invasion through the interaction with numerous signaling proteins such as Rac Family Small GTPase 1 (Rac1), p21-activated kinase 1 (PAK1), LIM domain kinase 1 (LIMK1) and Liver Kinase B1 (LKB1) (26–30). In ovarian cancer, increased expression of NISCH was associated with a block in the G1 phase of the cell cycle and thus a decrease in cell proliferation (31). Its down-regulation led to increased proliferation, migration and invasion of lung adenocarcinoma cancer cells (32). Our group has recently examined NISCH expression across human cancers and found that it was decreased in tumor tissues compared to their healthy counterparts in most solid tumor types, and – based on the gene set enrichment analysis – that it was negatively associated with pathways that control cancer growth and progression (33).

Of importance, there are several Food and Drug Administration (FDA) approved drugs that bind to IRAS-I1/NISCH (34), out of which the antihypertensive rilmenidine has been reported as the most potent agonist (35). Nischarin expression and its role in pancreatic cancer have not been investigated to date. Given the tumor suppressive properties of NISCH, we hypothesized that its agonist rilmenidine may have an effect on PDAC tumor growth and/or invasion and that it has a potential for drug repurposing in PDAC treatment. Here we present the findings on the anti-metastatic effects of rilmenidine on PDAC cancer cells and their interaction with the microenvironment.

## MATERIAL AND METHODS

### Acquisition and processing of data on NISCH expression from public repositories

The Cancer Genome Atlas (TCGA) pancreatic adenocarcinoma dataset was downloaded from the web-based bioinformatics tool cBioPortal (https://www.cbioportal.org/study/summary?id=paad_tcga_pan_can_atlas_2018) and merged with TCGA PAAD data from Morpheus web-site (https://software.broadinstitute.org/morpheus/) to obtain transcriptomic values of normal samples (36–38). PDAC datasets GSE16515 (39) and GSE62452 (40) from the NCBI Gene Expression Omnibus (GEO) database were downloaded in pre-processed form (41–43). To eliminate multiple probes for the NISCH gene, GEO datasets were uploaded to the Gene Set Enrichment Analysis (GSEA) Broad Institute software (44,45) to collapse dataset from probes to symbols using the max_probe collapsing mode. Raw RNA-seq results of GSE93326 data set (46) were downloaded from GEO database and pre-processed in R.

Sample values were then divided into different groups to determine the effects of patients’ sex, tumor stage and grade, and 3p status on NISCH expression in PDAC samples using t-test, one-way ANOVA and Dunnett’s Tukey’s multiple comparisons test in GraphPad Prism 7. NISCH expression in normal pancreas and PDAC samples was analyzed with paired t-test in two datasets - GSE16515 and GSE62452, that had matching normal and tumor samples from the same patients. Matching stromal and epithelial samples from GSE93326 were analyzed using the same method. Patients from databases containing information about survival (TCGA PAAD and GSE62452) were divided into the NISCH high and NISCH low expression groups based on the mean of NISCH expression of the entire population, and survival analysis was performed using the log-rank (Mantel-Cox) test.

The HPA database (https://www.proteinatlas.org/ENSG00000010322-NISCH/pathology/pancreatic+cancer, (47) was used to download images of tumor tissues stained with HPA023189 antibody (accessed on July 2^nd^ 2024).

GEPIA (Gene Expression Profiling Interactive Analysis, http://gepia.cancer-pku.cn/) (48) was used to examine the mRNA expression of NISCH in different stages of PDAC and generate box plots.

The expression level of total NISCH protein in normal pancreatic tissue and pancreatic adenocarcinoma from the Clinical Proteomic Tumor Analysis Consortium (CPTAC) was graphed with Jitter plot using UALCAN web tool (49,50), where Z-values represent standard deviations from the median across samples for the given cancer type and where Log2 Spectral count ratio values from CPTAC were first normalized within each sample profile, then normalized across samples (accessed on 30^th^ July 2024).

For infiltration estimation for PAAD TCGA tumors, TIMER2.0 website (http://timer.cistrome.org/) was used (51–53). We used Gene module to visualize the correlation of NISCH expression with immune infiltration level of B cells, CD8+, CD4+, dendritic cells and macrophages and neutrophils.

### Immunohistochemistry

The pancreas tumor tissue array was purchased from Novus Biologicals (NBP2-78128, Littleton, Colorado, USA). The tissue array included formalin-fixed duplicated 24 cases of normal, reactive and cancerous tissues of the pancreas. The array was deparaffinized by heating at 58°C for 1 h, followed by series of xylene washes, and hydrated in graded ethanol series. Antigen retrieval was performed in Abcam solution (ab208572) for 20 minutes in a 95°C water bath. After hydrogen peroxide block and rinse, subsequent reagents were from the Immunoperoxidase Secondary Detection System (DAB500, Merck Millipore, USA) and were applied according to manufacturer’s recommendations. Primary anti-nischarin mouse monoclonal antibody (BD Pharmingen, UK #558262) was used at 1:300 dilution and incubated at 4°C in humified chamber, overnight.

For immunofluorescent staining, primary antibody incubation with mouse anti-nischarin antibody in combination with rabbit anti-α-Smooth Muscle Actin (D4K9N) XP monoclonal antibody (dilution 1:150, 19245, Cell signaling technology, USA) or rabbit anti-FAP (E1V9V) monoclonal antibody (dilution 1:50, 66562, Cell signaling technology, USA) was performed at 4°C in humified chamber, overnight. For NISCH and CD45 co-staining combination of anti-nischarin rabbit antibody (1:30 dilution) (HPA023189, Atlas antibodies) and mouse monoclonal anti-CD45 antibody (1:100 dilution) was used (CD45-2B11, Invitrogen) in Diluent Booster solution (AR1016, Booster Biological Technology). After a series of washes slides were incubated with secondary antibodies: Goat anti-Rabbit IgG (H+L) Cyanine3 (dilution 2 μg/mL, A10520, Invitrogen, Carlsbad, California) and goat anti-mouse Alexa Fluor 488 secondary antibody (37120, Invitrogen, Carlsbad, California) at RT for 1h. Nuclei were stained with DAPI (dilution 1:1000, 90229, Merck Millipore, USA) for 5 minutes and slides were mounted with Antifade mounting medium (AR0036, Booster Biological Technology, Pleasanton, California, USA). Images were acquired on Zeiss PALM MicroBeam Axio Observer Z1 microscope (Carl Zeiss MicroImaging GmbH, Germany) with Zeiss LD Plan*-* Neofluar 20x */*0.4 Korr, ∞/0-1.5 mm lens and AxioVision software.

### Cell lines and culture conditions

The PANC-1 and MIA PaCa-2 cell lines were procured through inter-laboratory exchange with The Vinča Institute of Nuclear Sciences, originally obtained from the American Type Culture Collection, (ATCC, Manassas, Virginia, United States). Pancreatic Cancer Panel ATCC-TCP-1026 was purchased from the ATCC. Cells were grown in the media recommended by the ATCC. All culture media were supplemented with 10% fetal bovine serum (FBS) (Sigma-Aldrich, USA) and antibiotics solution (PS-B, Capricorn Scientific, Germany). All the cell lines were occasionally tested for the presence of mycoplasma by PCR(54), and if required, treated with MycoXpert (Capricorn Scientific, Ebsdorfergrund, Germany).

Primary human CAFs were propagated from three different patients (labeled as CAF1, CAF2, and CAF3, all derived from female patients aged 63-74 years, therapy naïve). Based on the national health laws and good clinical practice, the Ethical committee of the Clinical Center of Serbia approved patient tissue use under Protocol number 570/9, from 31.07.2020. All patients signed the informed consent form. CAFs were isolated by the outgrowth method described by Bachem et al (55). Briefly, tumor samples after pancreatoduodenectomy were immersed in saline and brought on ice from the clinic to the laboratory. Fragments were cut into 2–3 mm^2^ pieces, placed in a 6-well plate, and allowed to attach in a humidified 5% CO_2_ incubator for 30 minutes. DMEM (Sigma Aldrich, St. Louis, USA) with 4.5 g/L glucose, 10% FBS, and antibiotic-antimycotic (P06-07300, PAN Biotech, Germany) was gently added and the explants were observed over time under the microscope for cell outgrowth and proliferation. When the cells were 70% confluent around the explants, explants were removed and CAFs were dissociated from the epithelial cells by differential trypsinization. Cultures were tested for the absence of E cadherin and Cytokeratin 19. CAFs were not immortalized and spontaneously reached senescence around the 25th passage, which limited the number of assays we were able to perform on each CAF line (**Supplementary Table S1** lists the assays performed on CAF cell lines).

### Immunoblotting

Cellular proteins were extracted using Pierce RIPA cell lysis buffer (89901, ThermoFisher Scientific, USA) with the addition of phosphatase (4906837001, Roche, Basel, Switzerland) and protease (5892791001, Roche, Basel, Switzerland) inhibitors. Protein concentration was measured using a BCA protein assay kit (23225, Pierce BCA protein assay kit, Thermo Scientific, USA), and 20-30 μg of lysates were separated on 8% SDS-PAG electrophoresis gels. Proteins were transferred onto PVDF membrane (88518, Thermofisher Scientific, USA) and blocked with 5% BSA (BSA, 05479-250G, SIGMA, USA) or 5% skimmed milk (70166, Sigma-Aldrich, USA) in TBST buffer (20mM Trizma base pH 7.6, 137mM NaCl and 0.1% Tween-20 (P7949-100ML, Sigma-Aldrich, USA)), depending on the primary antibody. Primary antibodies were: anti-nischarin (clone D6T4X, dilution 1:1000, 85124, Cell signaling, USA), anti-fibronectin 1 (FN, Clone 10, dilution 1:10000, 610077, BD Biosciences, USA), anti-FAP (clone E1V9V, dilution 1:1000, 66562, Cell signaling, USA), anti-αSMA (clone D4K9N, dilution 1:1000, 19245, Cell signaling, USA), anti-β-actin (clone 8H10D10, dilution 1:2000, 3700, Cell signaling, USA), anti-gapdh (clone G-9, dilution 1:1000, sc-365062, Santa Cruz Biotechnology, Santa Cruz, California, USA), anti-MMP-9 (clone 36020, dilution 1:500, MAB936, R&D Systems, Minneapolis, Minnesota,USA) and anti-vinculin (clone 7F9, dilution 1:1000, 90227, EMD Milipore, Darmstadt, Germany). Membranes were incubated with primary antibodies at 4°C overnight. Horseradish peroxidase (HRP)-linked goat anti-mouse IgG secondary antibody (#7076, Cell signaling, USA) or HRP-linked goat anti-rabbit IgG secondary antibody (#7074, Cell signaling, USA), were used at 1:2000 dilution and membranes were incubated for 1h at RT. Signal was developed with SuperSignal™ West Pico PLUS Chemiluminescent Substrate kit (34580, ThermoFisher Scientific, USA) and imaged on Azure600 Blot Imaging System (AZI600-01, Azure Biosystems, Dublin, California, USA).

### Drug treatments and MTT viability assay

Rilmenidine hemifumarate (R-134), moxonidine hydrochloride (M1559), and clonidine hydrochloride (C7897) were procured from the Sigma-Aldrich, USA and dissolved in DMSO (dimethyl sulfoxide) according to manufacturer’s instructions.

For the MTT (3-(4,5-dimethylthiazol-2-yl)-2,5-diphenyltetrazolium bromide, Sigma Aldrich, USA) viability assay cells were seeded in the 96-well plates, at a density of 3 x 10^3^ cells/well (MIA PaCa-2), 4 x 10^3^ cells/well (PANC-1), 6 x 10^3^cells/well (AsPC-1, BxPC-3, CFPAC-1), and 1 x 10^4^ cells/well (HPAF-II, SW1990), and treated at the concentration range from 0 to 600 μM for 72 hours. Experiment was performed in triplicates, with at least three biological repeats. Absorbance of the formazan solution was measured at 570 nm (Scientific Multiskan SkyHigh Microplate Spectrophotometer, Thermofischer Scientific, Massachusetts, USA), and IC_50_ values (50% decrease in MTT reduction compared to untreated control) were determined from the dose-response curves in GraphPad Prism 8.0.1 (GraphPad Software Inc., California, USA).

### Apoptosis assay

PANC-1 and MIA PaCa-2 cells were seeded at 1.5 x 10^5^ cells/well, and BxPC-3 at 3 x 10^5^ cells/well in 6-well plates. At the 24 hours after seeding, the cells were treated with 100 μM or 300 μM rilmenidine and incubated with the drug for additional 24, 48, or 72 hours. Cell death was analyzed using an Annexin V-fluorescein isothiocyanate apoptosis detection kit, according to the manufacturer’s instructions (556419, BD Biosciences, Franklin Lakes, USA) on FACS-Calibur cytometer with Cell Quest computer software (Becton Dickinson, Heidelberg, Germany). Results are shown as % of viable (Annexin V^-^/PI^-^) cells using GraphPad Prism 8.0.1 (GraphPad Software Inc., California, USA).

### RNA sequencing and Gene ontology (GO) enrichment analysis

Total RNA was isolated using TRIzol reagent (15596018, Life Technologies, USA) and RNA concentration was measured using BioSpec-nano Spectrophotometer (Shimadzu Biotech, Japan). Samples were prepared in the final concentration of 20 ng/μL in water, and sent for sequencing at Novogene Co (Cambridge, UK).

Gene Ontology (GO) enrichment analysis of differentially expressed genes and graphic representation was provided by Novogene. It was implemented by the cluster Profiler R package, in which gene length bias was corrected, and GO terms with a corrected p-value of less than 0.05 were considered to be significantly enriched. Differential expression analysis of two conditions/groups (two biological replicates per condition) was performed using the DESeq2 R package (1.20.0). The resulting p-values were adjusted using the Benjamini and Hochberg’s approach for controlling the false discovery rate. Genes with the adjusted p-value ≤ 0.05 found by DESeq2 were assigned as differentially expressed. The genes were ranked according to the degree of differential expression in the two samples, and then the predefined Gene Set were tested to see if they were enriched at the top or bottom of the list.

### Cell Spreading Assay

PANC-1, MIA PaCa-2, and BxPC-3 cells were seeded at 5 x 10^3^ cells/well in 48-well plates (Costar, Corning), uncoated or pre-coated with 2 μg/cm^2^ fibronectin (F4759, Sigma-Aldrich, USA) or collagen type I (C3867, Sigma-Aldrich, USA). Cells were treated with 50, 100, or 300 μM rilmenidine and imaged at 4, 6, 8, and 24 hours post seeding. Phase-bright round cells were counted as non-spread; cells with any protrusions were counted as spread. Experiments were performed with at least three biological repeats, and results were expressed as the percent of spread cells out of the total counted per field.

### Wound-healing assay

PANC-1, MIA PaCa-2, and BxPC-3 cells were seeded in 6-well plates at 5 x 10^5^ cells/well in growth media supplemented with 10% fetal calf serum (FCS) (Sigma-Aldrich, USA). 24 hours after seeding, the wound was made with a 200 μL pipette tip, and detached cells were washed with PBS solution. To decrease the proliferation of cells and ensure that the closing of the wound was a consequence of the directional migration of cells, experiment was performed in a medium supplemented with 1% FCS. Images of at least two areas per well were taken at 0 and 24 hours after 100 or 300 μM rilmenidine treatment at 10X magnification (Olympus CX33, Germany). The closure area of the migrating cells was calculated using ImageJ software (1.53e, National Institutes of Health, USA), and the percentage of wound closure was calculated at 24 hours post wounding, normalized to untreated control.

### Transwell assays

For transwell migration, cells were seeded on 0.8 μm pore inserts (353097, Corning Scientific, Germany) in 24-well plates at a density of 5 x 10^4^ cells/insert. Cells were seeded in the upper chamber in the presence of rilmenidine or without it for the control sample, in a serum-free medium. The lower chamber medium supplemented with 10% FCS was used as a chemoattractant. Cells were incubated for 48 hours in a humidified atmosphere containing 5% CO_2_ at 37 °C. For transwell invasion assay, inserts were pre-coated with 2 µg/cm^2^ fibronectin (F4759, Sigma-Aldrich, USA), 10 µg/mL collagen type I (C3867, Sigma-Aldrich, USA) or 0.4-0.6 mg/mL Cultrex® Reduced Growth Factor Basement Membrane Extract (RGF-BME, 3433-005-01, R&D Systems, Bio-Techne, USA). Cells were seeded in the presence of rilmenidine or without it, as in migration assays, and left to invade for 72 hours. After the incubation, the media was decanted and the remaining cells in the upper chamber were removed using a cotton swab. Cells that migrated/invaded through pores were fixed with 3.7% formaldehyde for 15 minutes, and stained with 0.5% Crystal Violet Dye in 25% methanol, for 30 minutes. Images of migrated/invaded cells were taken at 10X magnification (Olympus CX33, Germany), in at least three independent fields of view. Experiments were repeated three times and an average number of invaded cells was expressed as a number of invaded cells or % of area of invaded cells relative to uncoated untreated control using ImageJ software.

### Gelatin zymography

PANC-1, MIA PaCa-2, and BxPC-3 cells (2 x 10^6^ /tissue culture dish) were seeded in the presence of rilmenidine in serum-free media and were incubated for another 6, 24, and 48 hours. After the incubation, conditioned media from the cell culture were collected and further concentrated by using the protein concentrator spin columns (88517, Thermo Scientific, USA). Cells were lysed by scraping on ice with 200 µL of Cell Lysis Buffer (K490-100-2, BioVision, UK), centrifuged, and the supernatant was used. Protein quantification was performed by the BCA method (23225, Pierce BCA protein assay kit, Thermo Scientific, USA). Gelatin zymography of conditioned media samples and cell lysates was performed on 8% SDS-PAGE containing 1 mg/mL of gelatin (48723, Fluka Analytical, USA) under non-reducing conditions, as previously described (56). After rinsing in renaturing buffer (2.5% Triton X-100), the gels were soaked in the developing buffer containing 50 mM Tris-HCl, 200 mM NaCl, 5 mM CaCl_2_, and 0.05% NaN_3_, pH 7.5 at 37°C for the next 48 hours. Gels were then stained with 0.5% Coomassie brilliant blue for 1 hour and destained in 40% methanol and 10% acetic acid. An additional destaining step with 1% Triton X-100 was performed to obtain sharper MMP bands over the blue background. The transparent bands corresponding to MMP activity were analyzed by optical densitometry with ImageJ software.

### Immunofluorescence staining

CAFs were seeded in 96-well optical plates (165305, Thermo Scientific, USA) at a density of 2 x 10^3^ cells/well. Cells were treated with 100 μM rilmenidine 24 hours after seeding and incubated for another 72 hours, after which they were fixed with 3.7% formaldehyde for 15 minutes. After the PBS wash, cells were permeabilized with 0.2% Triton X-100 (T-6878, SIGMA, USA) in PBS on ice for 2 minutes. Unspecific binding sites were blocked with 1.5% BSA in PBS for 30 minutes at room temperature. Fixed cells were incubated in primary antibodies: mouse anti-NISCH (dilution 1:150, 558262, BD Pharmingen, UK, rabbit anti-α-Smooth Muscle Actin (D4K9N) XP monoclonal antibody (dilution 1:150, 19245, Cell signaling technology, USA), rabbit anti-FAP (E1V9V) monoclonal antibody (dilution 1:50, 66562, Cell signaling technology, USA), anti-Pro Collagen I alpha 1 mouse monoclonal antibody (COL1α1, dilution 1:50, Clone # 816161, MAB6220, R&D Systems, Minneapolis, Minnesota, USA) and mouse anti-fibronectin (FN1, dilution 2.5 μg/mL, Clone 10, 610077, BD Biosciences, East Rutherford, New Jersey, USA), overnight at 4°C. After washing with PBS, secondary antibodies were added at 1:250 dilution: goat anti-mouse Alexa Fluor 488 (37120, Invitrogen, Carlsbad, California, USA, and goat anti-rabbit Alexa Fluor 488 secondary antibody (dilution 4μg/mL, A11008, Invitrogen, Carlsbad, California, USA) and incubated for 1 h at room temperature. Actin fibers were stained with Phalloidin TRITC Dye (dilution 1:500, 90228, Merck Millipore, Darmstadt, Germany) for an hour, and nuclei were stained with DAPI fluorescent dye (dilution 1:1000, 90229, Merck Millipore, USA) for 2 minutes. After washing with PBS, cells were imaged on a fluorescent microscope Zeiss PALM MicroBeam Axio Observer Z1 (Carl Zeiss MicroImaging GmbH, Germany) using 10X, 20X or 63X Zeiss LD Plan*-*Neofluar objectives.

### Co-culture of pancreatic cancer cells and CAFs

CAFs were seeded in 6-well plates at 5 x 10^5^ cells/well in DMEM supplemented with 10% FCS and Penicillin-Streptomycin solution. Inserts with 0.4μm pore size (CC Insert MD6 140640, Thermofisher Scientific, USA) were placed in wells, and PANC-1 cells were seeded into the inserts at the density of 3 x 10^5^ cells/insert. After 24 hours, the medium was replaced in both compartments with DMEM without serum or with DMEM without serum and 100 μM rilmenidine and co-cultures were incubated for another 72 hours. After incubation conditioned medium was collected for dot-blot and ELISA analyses and pellets of cells were collected for RT-PCR analysis.

### Dot-blot analysis

Conditioned media from the rilmenidine-treated and untreated co-cultures of PANC-1 cells with CAF1 or CAF2 cells were collected. The human cytokine array (ARY005B, R&D Systems, Minneapolis, Minnesota, USA) was used to determine differences in cytokine expression between treated and untreated samples. The experiment was performed according to the manufacturer’s instructions. Briefly, conditioned media was first concentrated on Amicon® Ultra-4 Centrifugal Filter Units (UFC800308, Merck Millipore, Darmstadt, Germany) for 20 minutes and total protein concentration was determined using the BCA assay (23225, Pierce BCA protein assay kit, ThermoScientific, USA). The volumes of samples used in the reaction were 400 μL for media from PANC-1 – CAF1 co-culture condition and 200 μL for media from PANC-1 – CAF2 co-culture condition. Human Array Detection Antibody Cocktail (15 μL) was added to dot-blot membranes and incubated overnight at 4°C. After washing, membranes were incubated in Streptavidin-HRP conjugate for 30 minutes at room temperature. For the detection, Chemi Reagent Mix was used and images were acquired on the Azure600 Blot Imaging System (AZI600-01, Azure Biosystems, Dublin, California, USA). The integrated intensity of the dots was determined using Image J software and normalized to the amount of the loaded protein. Cytokines for which the difference in amount was observed were plotted in GraphPad Prism 8.0.1 as integrated density per μg of protein.

### Enzyme-linked immunosorbent assay

Conditioned media from co-cultures of PANC-1 or MIA PaCa-2 cells and CAF2 cells were collected as described above. The concentration of total proteins was determined in the BCA assay. Interleukin-6, interleukin-8, and PAI-1 levels were detected using a commercial Human IL-6 ELISA kit (88-7066-22, Invitrogen, Carlsbad, California, USA), Human IL-8 ELISA kit (88-8086, Invitrogen, Carlsbad, California, USA) and Human SERPINE1 (Plasminogen activator inhibitor 1) ELISA kit (EH0538, Finetest, Wuhan, Hubei, China), respectively. ELISAs were performed according to the manufacturer’s instructions. Absorbance was measured at 450 nm using a microplate reader (Scientific Multiskan SkyHigh Microplate Spectrophotometer, Thermofischer Scientific, Massachusetts, USA). After measuring the absorbance, concentrations were determined based on the standard curve, and the results were normalized to the concentration of total loaded proteins.

### Analysis of gene expression by Real-time PCR

Total RNA was isolated using TRIzol reagent and was reversed transcribed into complementary DNA using a High Capacity cDNA RT kit (4368814, ThermoFisher Scientific, USA) as per manufacturer’s instructions. The qPCR was performed using SYBR Green Master Mix (A25742, Applied Biosystems, USA) and gene expression was calculated by the 2^-ΔΔCt^ method. Sequences of the primers used for analysis are shown in **Table 1**.

**Table 1.**
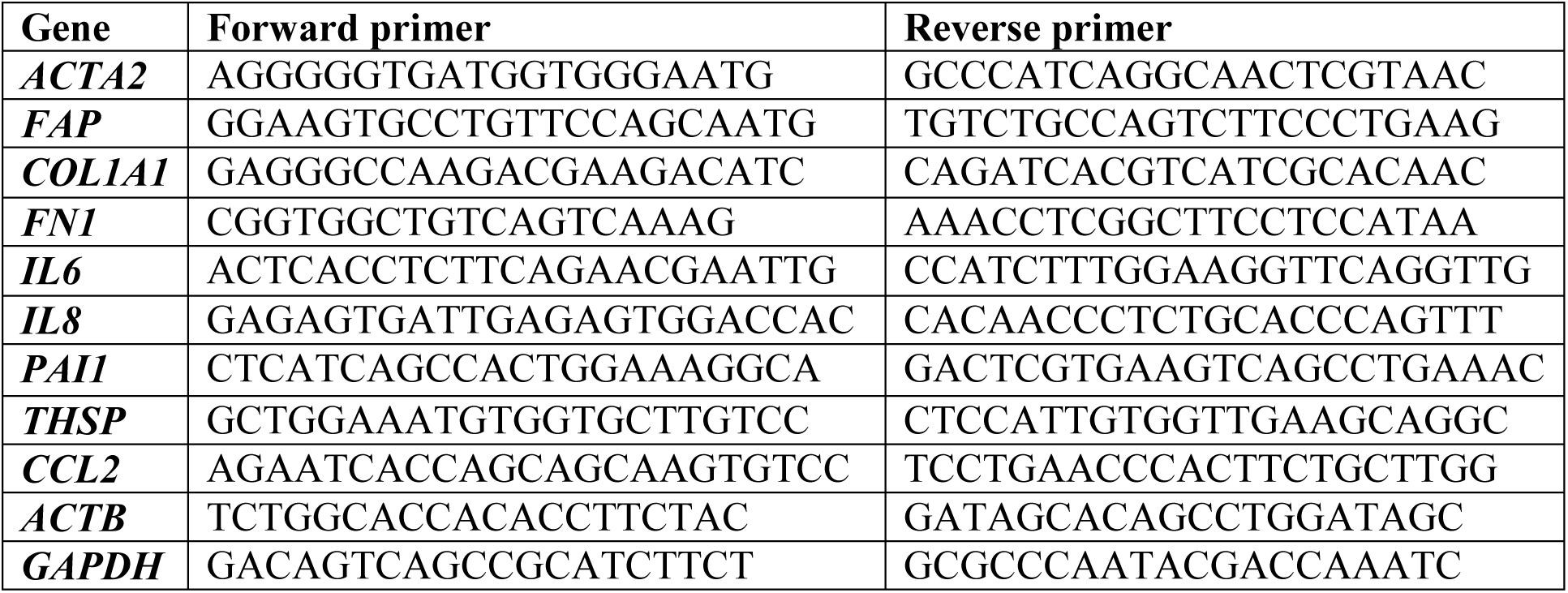
Sequences of the primers used for qRT-PCR analysis.

### Zebrafish pancreatic adenocarcinoma xenograft model

All procedures with zebrafish (*Danio rerio*) embryos complied with the European Directive 2010/63/EU and adhered to the Ethical Guidelines for the Care and Use of Laboratory Animals established by the Institute of Molecular Genetics and Genetic Engineering, University of Belgrade. Wild type zebrafish embryos (AB strain), kindly provided by Dr. Ana Cvejić (Wellcome Trust Sanger Institute, Cambridge, UK), were reared to adulthood in a temperature- and light-controlled facility at 28°C with a standard cycle of 14 hours of light and 10 hours of darkness. The adults were fed twice daily with commercial dry food (SDS200 and SDS300 granular diets from Special Diet Services, Essex, UK, and TetraMin™ flakes from Tetra Melle, Germany) and supplemented daily with *Artemia nauplii*.

The toxicity of rilmenidine was evaluated in accordance with the general principles of the OECD guidelines for chemical testing. Embryos resulting from pairwise mating were washed to remove debris and divided into 24-well plates, with 10 embryos per well in 1 mL of E3 medium (consisting of 5 mM NaCl, 0.17 mM KCl, 0.33 mM CaCl_2_ and 0.33 mM MgSO_2_ in distilled water). The plates were kept at 28°C throughout the experiment. To evaluate acute (lethal) and developmental (teratogenic) toxicity, zebrafish embryos were exposed to the five different concentrations (5, 12.5, 25, 50 and 100 µM) of rilmendine at the 6-hour post-fertilization (hpf) stage (to ensure high sensitivity to the applied molecules) and at the 48 hpf stage (corresponding to the start of treatment for the pancreatic zebrafish xenografts, see the following section). Rilmenidine, prepared as a DMSO 42 mM stock, was diluted directly in E3 medium for exposure. DMSO (0.25%) and E3 medium alone was used as negative control. Treated embryos were examined every day under a stereomicroscope (Carl Zeiss™ Stemi 508 doc, Germany) for the appearance of apical endpoints up to 120 hpf (**Supplementary Table S2**). Dead embryos were discarded every 24 hours. The experiment was performed in triplicate with 20 embryos per concentration. At 120 hpf, embryos were anesthetized by the addition of 0.1% (w/v) Tricaine solution (Sigma-Aldrich, St. Louis, MO), photographed, and euthanized by freezing at −20 °C for ≥ 24 h.

For the xenograft experiments PANC-1 cells were detached with trypsin, washed three times PBS and resuspended in serum-free DMEM medium. Prior to microinjection, the cells were labeled with 2 μM Red CellTracker fluorescent dye according to the manufacturer’s instructions and then washed with PBS. Approximately 5 nL of the suspension containing 150 labeled PANC-1 cells was microinjected into the yolk sac of transgenic Tg(*fli1*:EGFP) zebrafish embryos (with fluorescently labeled vasculature) at 48 hpf using a pneumatic picopump (PV820, World Precision Instruments, USA). The injected embryos were incubated at 28°C for one hour to recover and examined under a fluorescence microscope to confirm the presence of fluorescently labeled tumor cells. Four hours post microinjection, successfully xenografted embryos were treated with 50 µM or 100 µM rilmedine and maintained at 33°C until the 120 hpf stage (3 days of treatment). The control group comprised xenografted embryos treated with 0.25% DMSO. The treated embryos were examined daily by fluorescence microscopy to evaluate the effect of rilmenidine on the survival of PANC-1 xenografts, the number of xenografts with metastases and the metastatic cell dissemination rate. The experiment was repeated twice, with 10 embryos per treatment.

### Statistical analysis

Numbers of repeats of experiments are shown in figure legends and all results were expressed as mean ± SE or mean ± SD. Statistical analysis was performed using GraphPad Prism 8.0.1 (GraphPad Software Inc., California, USA). Differences between control and treatment were analyzed with paired or unpaired two-tailed Student’s t-test, one-way or two-way ANOVA with a p < 0.05 considered as statistically significant.

### Study approval

All human samples used in this study were obtained from patients with written informed consent. The study was approved by the Ethics Committee of the University Clinical Center of Serbia and the Ethics Committee of the Institute for oncology and radiology of Serbia (approval numbers 570/9, from 31.07.2020 and 01-1/2023/703 respectively).

### Data availability statement

The data that support the findings of this study are available from the corresponding author upon reasonable request. The public data of NISCH mRNA and protein expression levels in PDAC and normal pancreatic tissues in this study were obtained from public repositories https://portal.gdc.cancer.gov, https://www.cbioportal.org/, http://gepia2.cancer-pku.cn/, https://ualcan.path.uab.edu/ and www.proteinatlas.org/. Raw sequencing data of MiaPaca2 cell line mRNA transcriptome, treated for 24h or 48h *in vitro* with 100uM rilmenidine can be found at https://zenodo.org/records/6920520 and https://zenodo.org/records/6948536 respectively.

## RESULTS

### 1. Nischarin is expressed in PDAC and is a favorable prognostic marker

We first examined NISCH mRNA and protein expression in the publicly available databases. NISCH mRNA and protein were expressed both in the healthy and tumor pancreatic tissues in the HPA, and there was no difference in the mRNA levels between the healthy and cancer tissue neither in the TCGA PAAD dataset (**Fig. 1A**) nor in the paired tumor and adjacent tissue samples in the GSE62452 (40) (**Fig. 1B**) and GSE16515 (57) datasets (**Supplementary Fig. 1A**). In the TCGA PAAD cohort, tumors of stage IIa and IIb had lower *NISCH* mRNA expression than stage I tumors (**Fig. 1C**), while in the GSE62452 cohort only stage IV tumors had lower expression than stage I, by Tukey’s multiple comparisons test (**Fig. 1D**). There was no difference in expression based on the grade of the tumor, nor between male and female patients (**Supplementary Fig. 1B and C**). Using the mean expression cut-off for separation into the high and low *NISCH* expression groups, we found that *NISCH* was a positive prognostic marker in terms of both progression free and overall survival of PDAC patients in the TCGA PAAD cohort (p=0.01 and p=0.018, by Mantel-Cox test, respectively) (**Fig. 1E, F**). In the GSE62452 cohort patients with higher *NISCH* mRNA expression had better survival, although the difference did not reach significance (p=0.19 by Mantel-Cox test, **Fig. 1G**). *NISCH* gene is located on the 3p21.1 chromosome, location marked as a putative tumor suppressor cluster (58). In the TCGA PAAD dataset based on the 3p status, tumors with the 3p loss had significantly lower *NISCH* expression (**Fig. 1H**).

**Figure 1.**
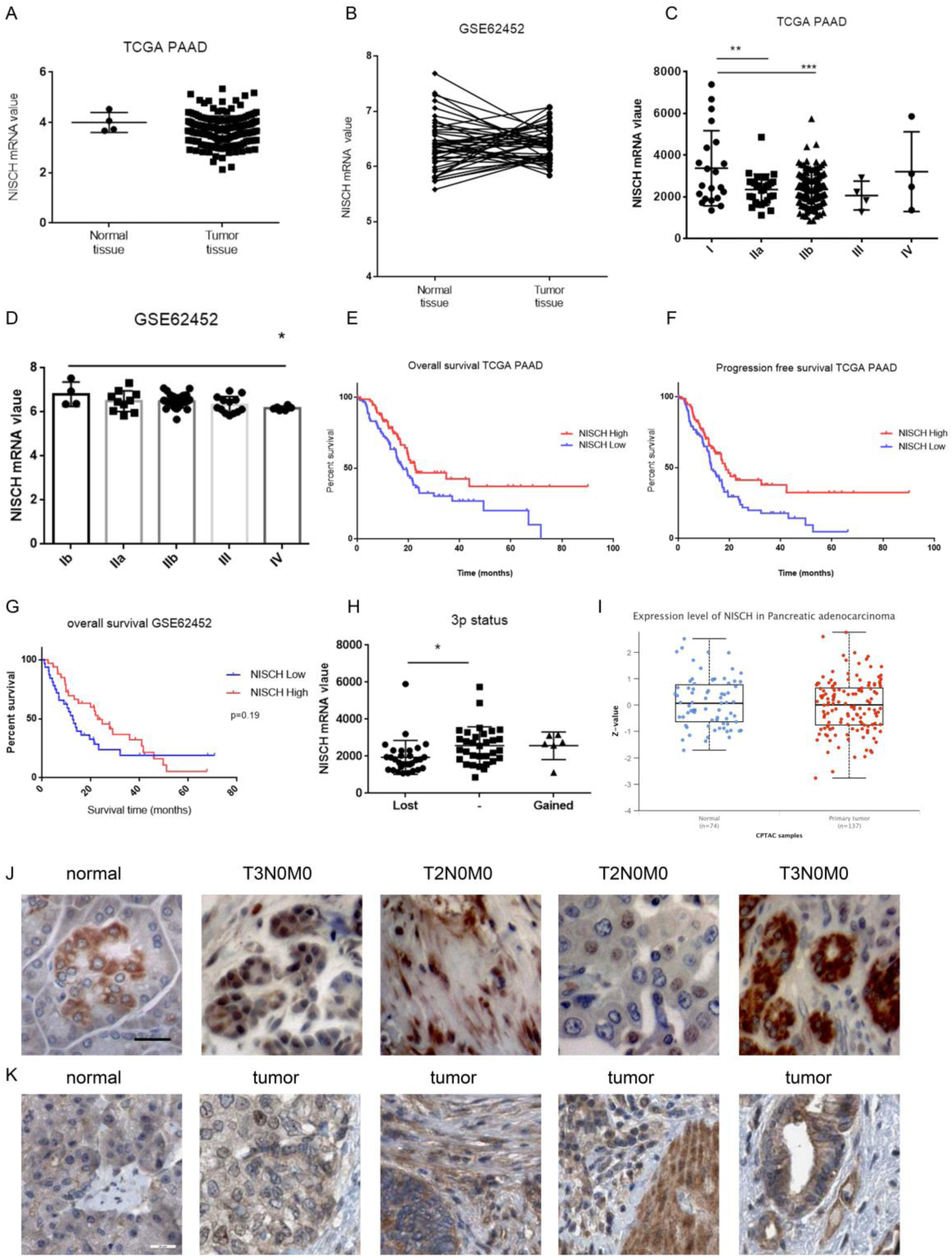
Nischarin is expressed in PDAC. A) *NISCH* mRNA expression in normal and tumor pancreatic tissues from the TCGA PAAD cohort; B) *NISCH* mRNA expression in the paired tumor and adjacent tissue samples in the GSE62452 cohort; C) *NISCH* mRNA expression by stage in the TCGA PAAD cohort, **p = 0.0052, ***p = 0.0003, adjusted p-value, one-way ANOVA, Tukey’s multiple comparison test, and D) the GSE62452 cohort, *p = 0.0414, one-way ANOVA, Tukey’s multiple comparison test; E) Kaplan-Meier plots for the overall and F) progression free survival of PDAC patients in the TCGA PAAD cohort divided into two groups using the best cut-off value for NISCH. p = 0.01 and p = 0.018 by Mantel-Cox test, respectively; G) The overall survival of patients in the GSE62452 cohort (p=0.19 by Mantel-Cox test); H) *NISCH* mRNA expression in groups based on the status of the 3p chromosome, in patients from the TCGA PAAD cohort, *p = 0.0299, one-way ANOVA, Tukey’s multiple comparison test; I) Total NISCH protein level in normal and cancer pancreatic tissue from CPTAC; J) NISCH expression in the normal and tumor pancreatic tissue from the NBP2-78128 microarray, scale bar 20 μm; K) NISCH expression in the normal and pancreatic tissue deposited in the Human Protein Atlas (HPA023198), scale bar 20 μm.

In the CPTAC database, mass-spectrometry-based proteomic data on NISCH protein expression was available on healthy and tumor tissue, but NISCH was not significantly lower in the tumor compared to the normal pancreas **(Fig 1I).** Next, we examined NISCH protein localization by immunohistochemistry (anti-nischarin #558262, BD Pharmingen) in the PDAC tissue microarray (NBP2-78128, Novus Biologicus). Nischarin was expressed in the normal pancreatic tissue, and in most of the examined pancreatic cancer tissue samples (**Fig. 1J**). Intriguingly, apart from the membranous and cytoplasmic localization, we observed NISCH in the nuclei of cells in tumor tissues. To further corroborate this, we examined NISCH expression and assigned localization in the HPA, in which samples were stained with a different antibody (HPA023198). Indeed, in the HPA, NISCH expression in PDAC was described as membranous, cytoplasmic and nuclear (in 50% of the samples) (**Fig. 1K**). Of importance, in the tissue microarray we examined, as well as in the HPA, NISCH could be observed in both the epithelial and the stromal compartments of the tumor.

### 2. NISCH is expressed in both the epithelial and stromal compartments of PDAC

Any systemic therapy for PDAC would potentially reach all of the components of the tumor tissue, both cancer and the stromal compartments. To infer *NISCH* mRNA localization, we examined the data from the GSE93326 dataset (46) that has paired laser-microdissected epithelial and stromal compartment samples from 65 patients. Intriguingly, *NISCH* mRNA expression was higher in the stromal than in the epithelial compartment (p=0.0001 by paired t-test) (**Fig. 2A**), but there was no difference in the expression between the ECM-rich and immune-rich stromal compartments (**Fig. 2B**). To validate this finding, we co-stained for NISCH with two markers for cancer-associated fibroblasts, fibroblast activation protein (FAP) and alpha smooth muscle actin (αSMA) (17), in the TMA. We found that NISCH was expressed in both FAP- (**Fig. 2C**) and αSMA-positive cells (**Fig. 2D**). In line with our findings, based on the Timer estimate (52,53,59) *NISCH* expression did not associate with tumor purity in PDAC, and partially correlated with various immune-cell infiltrates (**Supplementary Fig. 1D**). We stained for NISCH and CD45 in the TMA, but found very little CD45-positive infiltrate in the samples, apart from the liver metastasis (**Supplementary Fig. 1E**). Finally, we examined NISCH expression in a panel of human PDAC cell lines representing both classical and quasimesenchymal phenotype (MIA PaCa-2, PANC-1, Capan-2, BxPC-3, HPAF-II, CFPAC-1, AsPC-1 and SW 1990) and in CAFs isolated from the PDAC tumor tissue. NISCH was expressed in all the tested cancer cell lines (**Fig. 2E**) and in CAFs (**Fig. 2F**). These results confirm that NISCH is expressed in both epithelial and stromal compartments of PDAC tissue, and imply that any drug targeting NISCH would have an impact on both the cancer cells and the stroma.

**Figure 2.**
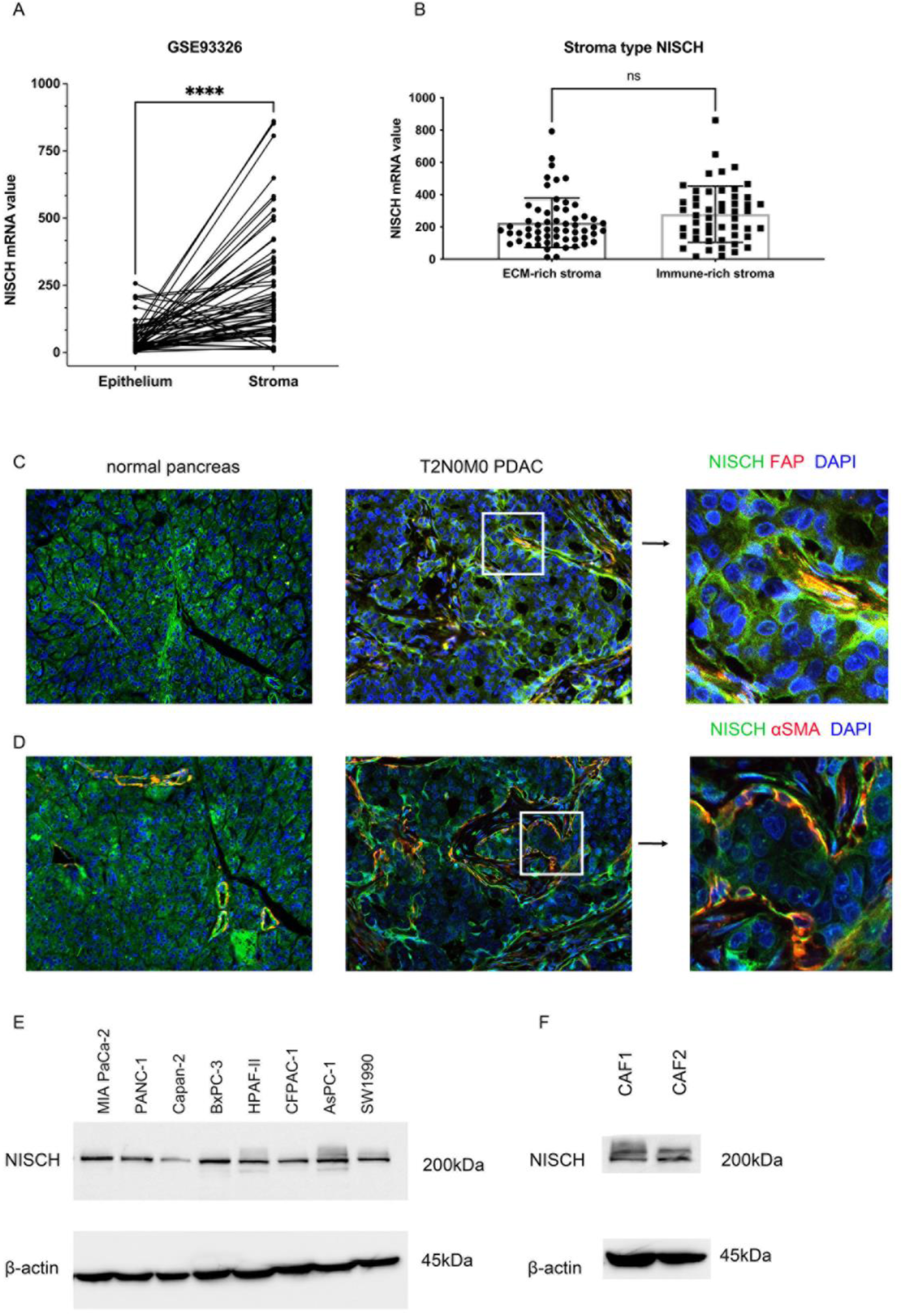
Nischarin is expressed in both the epithelial and stromal compartments of PDAC. A) *NISCH* mRNA expression in paired epithelial and stromal compartments of PDAC from the GSE93326 cohort, ****p < 0.0001, two-tailed paired t-test; B) *NISCH* mRNA expression in the ECM-rich and immune-rich type stroma in the GSE93326 cohort; C) NISCH (green) and FAP (red) localization in the normal and tumor pancreatic tissue from the NBP2-78128 microarray, nuclei blue; D) NISCH (green) and αSMA (red) localization in the normal and tumor pancreatic tissue from the NBP2-78128 microarray, nuclei blue; E) NISCH protein expression in PDAC cancer cell lines, and in F) patient-derived CAFs.

### 3. Nischarin agonists affect pancreatic cancer cell fitness and induce significant transcriptional changes *in vitro*

To examine the effects of I_1_-Imidazoline receptor IRAS-I1/NISCH agonists on cancer cell fitness, we chose the cell lines derived from the primary tumor site – MIA PaCa-2, PANC-1, Capan-2, and BxPC-3 – and tested the effects of rilmenidine and moxonidine (IRAS-I1 selective agonists from the second generation of central antihypertensive drugs) and clonidine (the first generation non-selective IRAS-I1 agonist). We performed the MTT viability assay after 72 hours of treatment with increasing drug concentrations (0 to 600 μM). We found that rilmenidine, with the highest affinity for NISCH out of the three tested drugs (34,60), had the most potent effect in all four tested cell lines, while clonidine had no effects on cell viability (**Fig. 3A**). The least sensitive to all three drugs was the Capan-2, cell line that expressed the lowest amount of NISCH. We selected rilmenidine as the most potent for further investigation and examined the sensitivity of the rest of the PDAC cell lines from the panel (IC_50_ values, **Supplementary Table S3**). MIA PaCa-2 cell line was the most sensitive to the treatment with the IC_50_ value of 169,2 μM. As the MTT test is based on the reduction of the formazan dye (61) and decreased viability observed in this test can be a consequence of the decreased proliferation, increased cell death, or the altered redox state in the cells, we tested the effects of 100 μM and 300 μM rilmenidine treatment on the apoptosis in the three primary tumor cell lines sensitive to rilmenidine: MIA PaCa-2, PANC-1, and BxPC-3. Flow cytometry analysis of the percentage of live cells negative for AnnexinV/PI staining showed a significant reduction of cell viability after 72 h of treatment with 300 μM rilmenidine on all three cell lines, with no significant changes under 100μM rilmenidine treatment at all time points (**Fig. 3B, C and D**). For the MIA PaCa-2 and PANC-1 cell lines, these results are in line with the MTT assay, while in the case of the BxPC-3, cell apoptosis and necrosis occur at much lower doses of rilmenidine than those predicted by the MTT assay. Taken together, rilmenidine treatment induces PDAC cell death only when administered at high concentrations. To infer other possible effects of NISCH agonization in PDAC cells, we sequenced the transcriptome of the MIA PaCa-2 cell line treated with 100 μM rilmenidine for 24 or 48 hours. Based on the Gene Ontology (GO) functional gene set enrichment analysis, 24 h treatment with rilmenidine had the most prominent negative effect on the transcripts associated with focal adhesion, adherens junction, cell-substrate adherence junction, positive regulation of cell migration and the extracellular matrix organization (**Fig. 3E**). These findings are in line with the so far reported tumor suppressive functions of the NISCH protein (62). After 48 h of treatment, in addition to the cell-substrate interactions and focal adhesion-associated transcripts, pathways associated with translation, ribosome assembly and protein targeting to various localizations in the cells were negatively associated with treatment (**Fig. 3F**). All these are related to proper cytoskeletal organization and imply that rilmenidine treatment affects the cytoskeletal dynamics.

**Figure 3.**
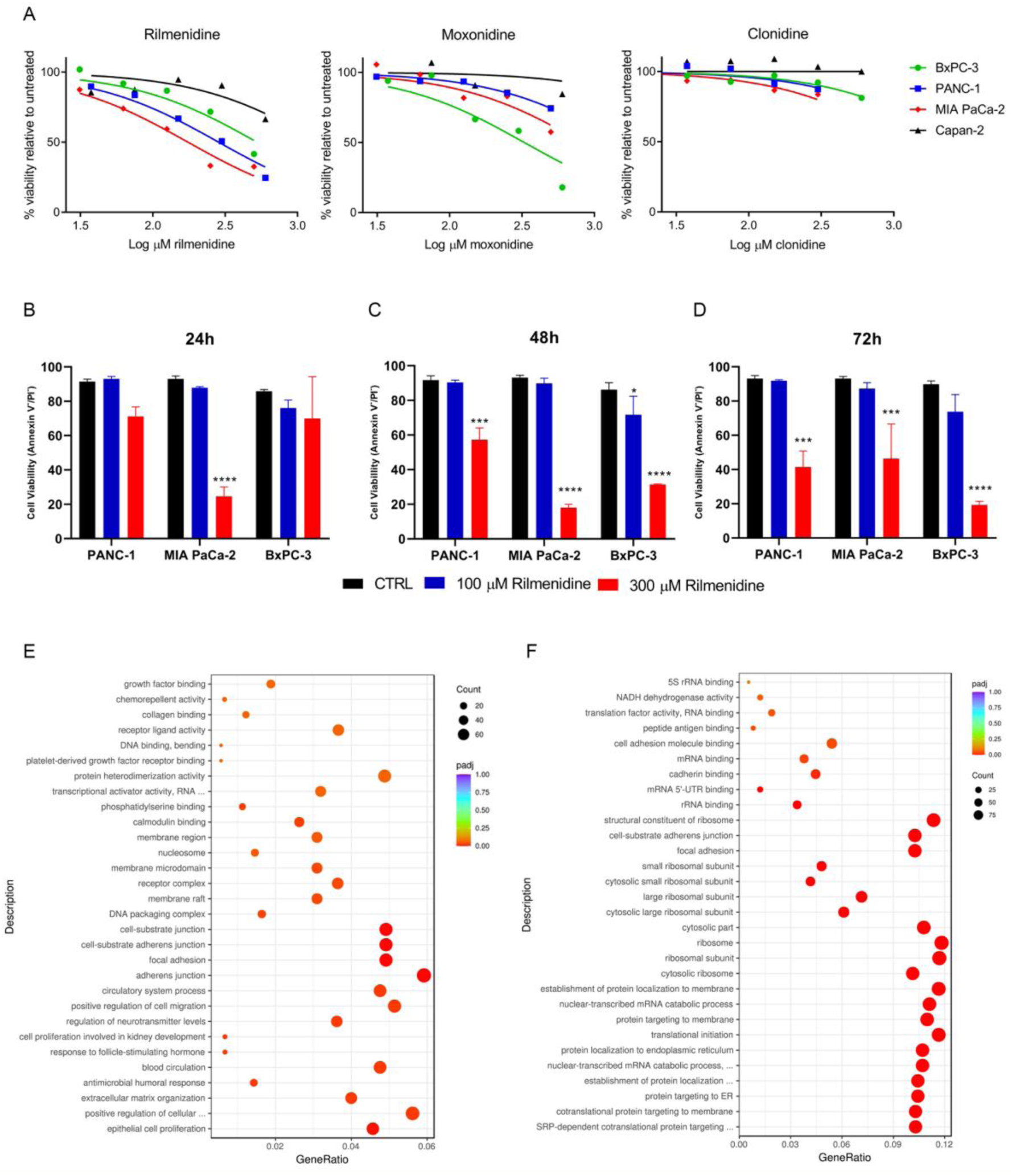
Effects of NISCH agonist treatment on PDAC cells *in vitro.* A) Percent viability of PDAC cells after 72 hours of treatment with increasing concentrations of rilmenidine, moxonidine or clonidine determined by MTT assay, presented as log(inhibitor) vs. normalized response; B) The proportion of live Annexin V^-^/ PI^-^ cells in the population of untreated and cells treated with 100 and 300 μM rilmenidine for 24 hours, C) 48 hours, or D) 72 hours. All data are shown as mean ± SD. n = 2; *p < 0.05, ***p < 0.001, ****p < 0.0001, two-way ANOVA (Sidak’s multiple comparisons test); E) Gene Ontology functional gene expression enrichment analysis of MIA PaCa-2 cells treated with 100 μM rilmenidine for 24 hours or F) 48 hours.

### 4. Nischarin agonist rilmenidine inhibits migration and metastatic spread of PDAC cancer cells

NISCH binds to integrin α5 and is involved in the regulation of focal adhesion assembly during cancer cell migration (24). Therefore, we first investigated the effect of rilmenidine on cell attachment on plastic, fibronectin- and collagen-coated surfaces. We found that rilmenidine delayed cell attachment in all the examined cell lines regardless of the surface coating, while 300 μM treatment completely prevented cell spreading (shown at 6 h and 24 h in **Fig. 4A**, at all**-**time points in **Supplementary Fig. S2A, B, C**, photomicrographs in **Supplementary Fig. S2D, E** and **F,** in the left panels). As expected, MIA PaCa-2 cell line had a delay in cell spreading on collagen I, given that it does not express integrin α2 and very poorly attaches to collagen-coated matrices (63). Next, we examined the effects of rilmenidine treatment on collective cell migration in wound-healing (WH) assay and chemotaxis in transwell (TW) migration assay. Rilmenidine treatment impaired cell migration in both WH and TW assays (**Fig. 4B, C**), with more potent effects in the TW assay where cells individually migrate. Representative images of the WH assay are shown in **Supplementary Fig. S2D, E** and **F,** in the right panels. To investigate the effects of rilmenidine treatment on cell invasion *in vitro,* we performed TW invasion assays with coating with three types of matrices: collagen I, fibronectin or matrigel (RGF-BME). 100 μM rilmenidine significantly reduced cell invasion only in the BxPC-3 cell line, regardless of the coated matrix (**Fig. 4D** quantification, representative images of TW invasion through matrigel coat **Fig. 4E)**. This result is not surprising, as PDAC cells *in vitro* are phenotypically plastic in terms of the migration mode and can adopt both mesenchymal (adhesion- and MMP-dependent) and amoeboidal (adhesion- and MMP-independent) migration modes (64). In the mesenchymal mode of invasion, cells degrade the ECM, partly through the secretion and/or activation of matrix metalloproteinases (MMPs) (65). We examined whether rilmenidine affects the activity of collagenases in conditioned media by gelatin zymography assay. Based on the predicted molecular size, we observed that BxPC-3 cells mostly secreted the MMP-9, while PANC-1 secreted mostly MMP- 2 (**Supplementary Fig. S3A and B**). MIA PaCa-2 cell line did not produce detectable MMPs under these culture conditions. Rilmenidine slightly reduced MMP gelatinolysis but the changes were not statistically significant (**Supplementary Fig. S3C**). Expression levels of MMP-9 in BxPC-3 cells increased over time (**Supplementary Fig. S3D**) but without significant changes upon rilmenidine treatment. Western blot of the conditioned medium of BxPC-3 cells confirmed a slight decrease of the MMP-9 expression after the rilmenidine treatment (**Supplementary Fig. S3E**). These results together imply that the effect of rilmenidine on cell migration and invasion are mostly the consequence of a decrease in cell adhesion properties under treatement.

**Figure 4.**
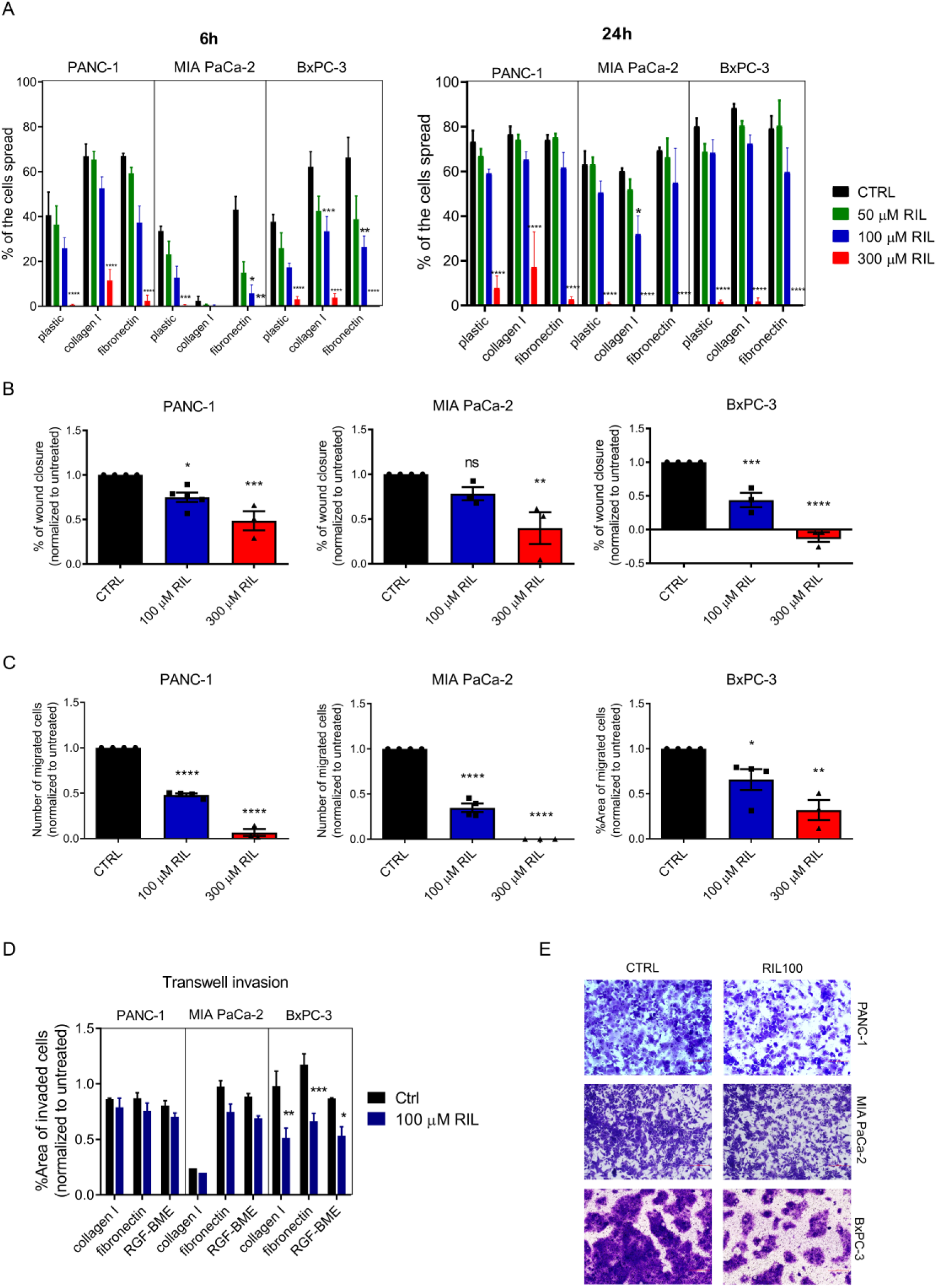
NISCH agonist rilmenidine reduces cell migration and invasion *in vitro*. A) Quantification of the number of PDAC cells spread 6 and 24 hours after seeding on tissue culture plastic, collagen I or fibronectin-coated surface in the presence of rilmenidine; B) Quantification of the PDAC cell migration in the wound healing assay after 24 h of treatment with rilmenidine; C) Quantification of the PDAC cell migration in the transwell assay after 48 h of treatment with rilmenidine; D) Quantification of PDAC cell invasion through collagen I, fibronectin or matrigel in the transwell assay after 72 h of treatment with rilmenidine; E) Representative images of transwell invasion through matrigel coat (right). All data are shown as mean ± SD. n = 3; *p < 0.05, **p < 0.01, ***p < 0.001, ****p < 0.0001, two-way ANOVA (Sidak’s multiple comparisons test).

Finally, we used the zebrafish xenograft model to assess the effects of rilmenidine on metastatic dissemination of tumor cells *in vivo*. This model leverages the optical transparency of zebrafish embryos, enabling real-time visualization of the metastatic spread of PANC-1 cells labeled with CellTracker Red through the body of transgenic *Tg*(*fli1*:EGFP) embryos with EGFP-expressing vasculature (**Fig. 5A**). To evaluate the potential therapeutic benefit of rilmendine in the treatment of pancreatic adenocarcinoma, we first investigated its dose-dependent toxicity *in vivo* using the zebrafish (*Danio rerio*) model. After continuous exposure for 5 consecutive days to various rilmendine’s doses ranging from 12.5 to 100 μM, the treated zebrafish embryos showed no visible signs of toxicity at any of the doses tested, including effects on survival, cardiac function, liver toxicity and skeletal development (teratogenicity).Over the course of a 3-day treatment, rilmenidine significantly inhibited the metastatic dissemination of pancreatic tumor cells throughout the vasculature and body of treated compared to untreated xenografts. Quantitative analysis revealed that rilmenidine significantly reduced both the proportion of embryos with metastases (at 100 µM, **Fig. 5B**) and the number of metastatic foci per embryo (at 50 µM and 100 µM, **Fig. 5C**). These findings demonstrate that the treatment with rilmenidine effectively reduces pancreatic cancer cell attachment and migration, leading to a substantial reduction in metastatic spread in *in vivo* xenograft model.

**Figure 5.**
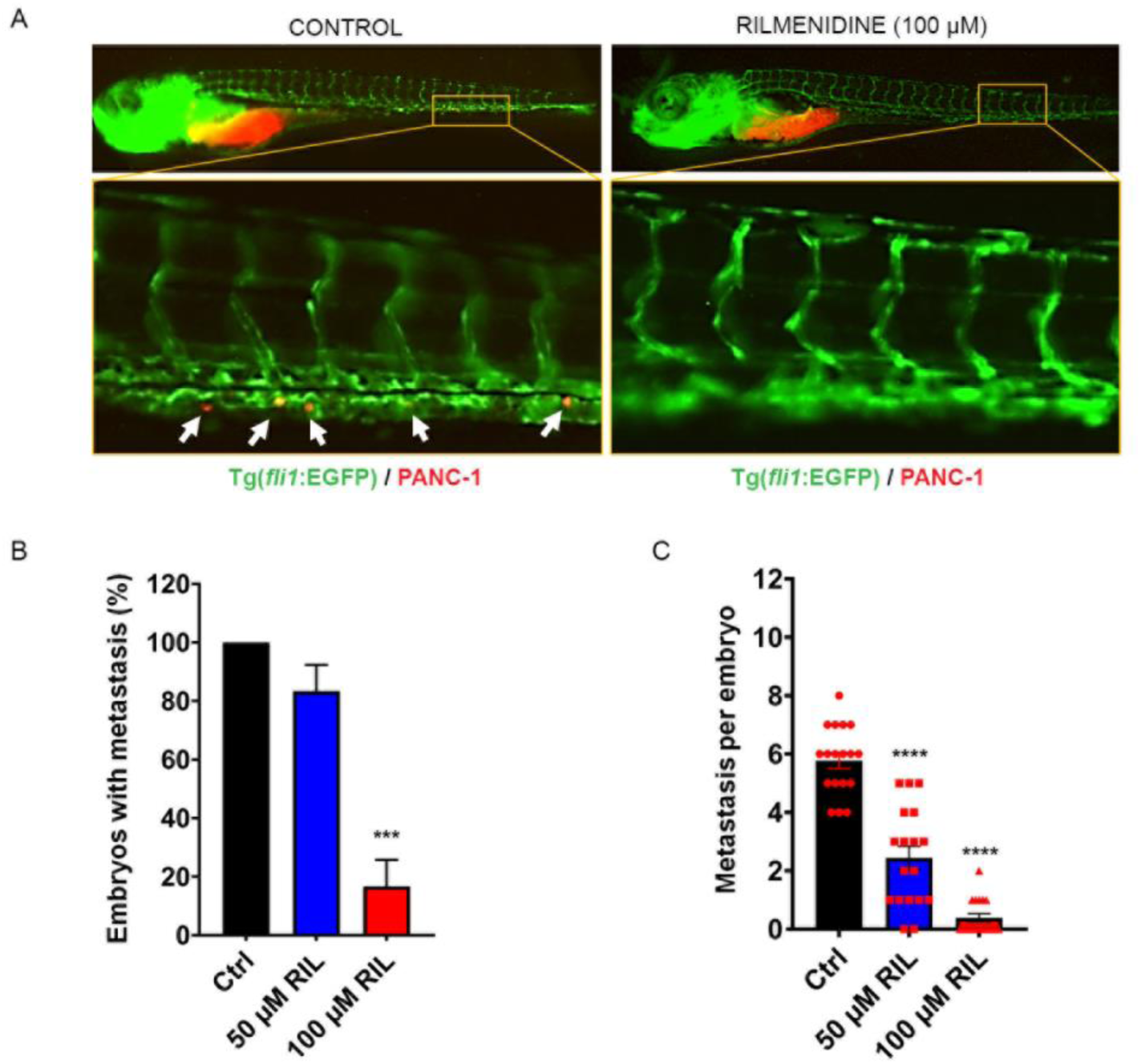
Rilmenidine reduces cell invasion in Tg(*fli1*:EGFP) zebrafish model. A) Effects of rilmenidine on PANC-1 cell growth and invasion in Tg(*fli1*:EGFP) zebrafish model. Squares amplify the indicated regions of metastatic sites. Arrowheads point to disseminated PANC-1 cells at 120 hpf. B) Quantification of the percent of embryos with metastasis and C) total numbers of metastatic foci in the caudal region of zebrafish. The graphs show mean ± SD, ***p < 0.001, ****p < 0.0001, One-way ANOVA (Dunnett’s multiple comparisons test).

### 5. Rilmenidine has an effect on CAF phenotype *in vitro* and modulates cancer cell-CAF communication

NISCH was also expressed in the stromal compartment of PDAC and in isolated CAFs from PDAC tumor tissue (**Fig. 2F**, **Fig. 6A**), therefore we examined the effects of rilmenidine on CAF cell fitness. In the MTT assay with up to 600 μM of rilmenidine treatment for 72 h, there were no effects on CAFs viability (**Fig. 6B**). Next, we examined the expression of CAF markers after treatment with 100 μM rilmenidine - FAP, αSMA, fibronectin (FN1) and collagen I alpha 1 (COL1α1). Immunofluorescence staining (**Fig. 6C, Supplementary Fig**. **S4A and B**) showed varied extent of decrease in expression of αSMA and deposition of, FN and COL1 that was cell line dependent, while FAP expression remained unchanged. Immunoblotting confirmed decreased expression of FN and αSMA in all three cell lines (**Fig. 6D and Supplementary Fig**. **S4C and D**). In the CAF2 line there was also a reduction in FAP expression upon treatment (**Fig. 6D**). As secreted growth factors in CAF-cancer cell co-culture support cell growth (66), we examined whether the observed changes in CAFs hold true in the presence of cancer cells. We cultured CAF2 or CAF3 cells in TW co-culture with PANC-1 or MIA PaCa-2 cells and examined CAF marker expression by immunoblotting. We observed a decrease in the expression of αSMA and FN in the presence of rilmenidine compared to control in the co-cultured cells, but also a decrease in the expression of FAP (**Fig. 6E, Supplementary Fig**. **S4D**). At the mRNA level, *αSMA, FAP* and *FN1* all decreased in the CAF2 monoculture upon rilmenidine treatment, and FN decrease was maintained in co-cultures with PANC-1 and MIA PaCa-2, while the effects on *αSMA* and *FAP* were cell line-dependent (**Fig. 6F**). Taken together, these results imply that rilmenidine may affect both the mRNA transcription and the protein stability of CAF markers.

**Figure 6.**
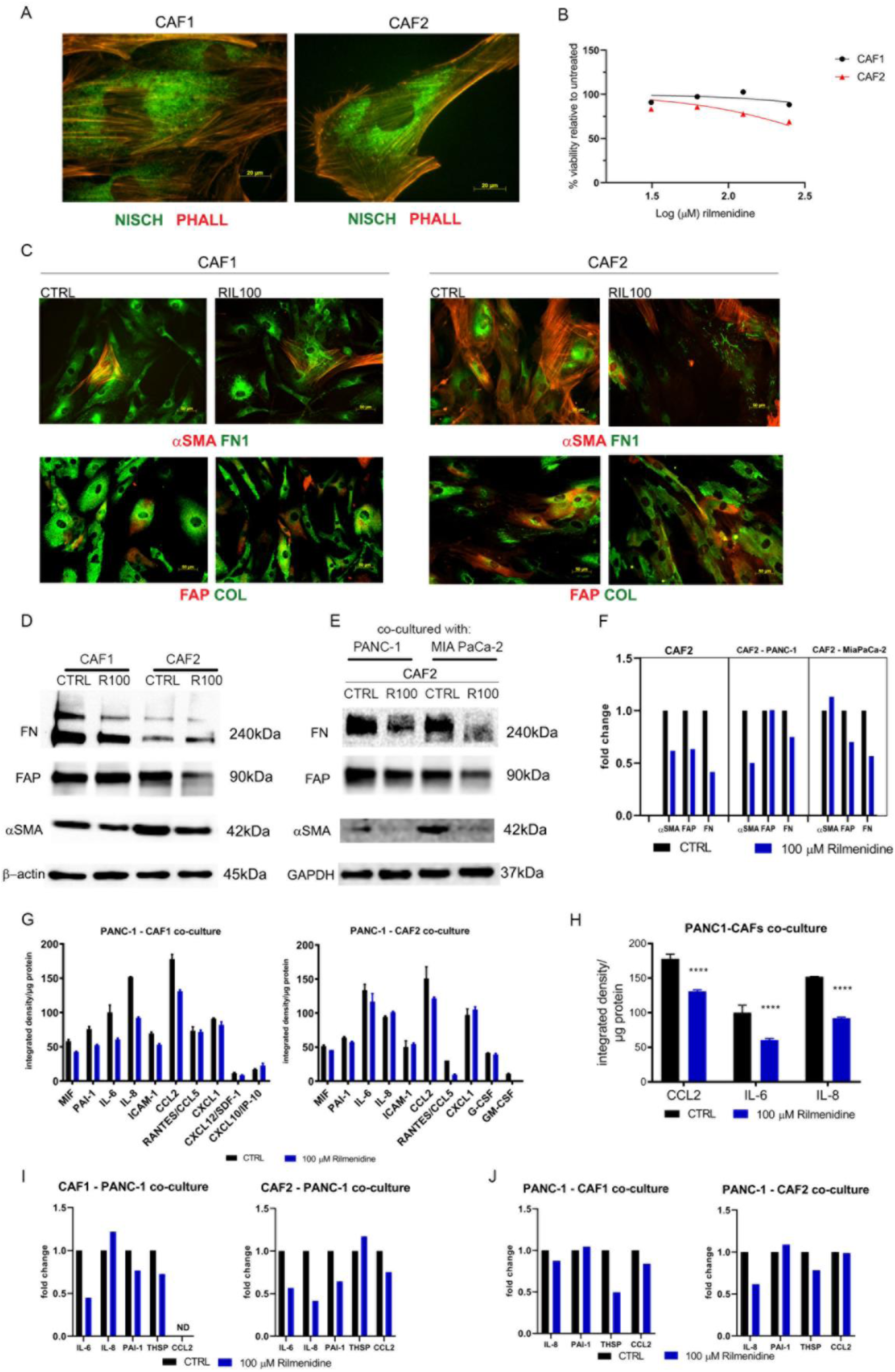
Rilmenidine affects the CAF phenotype and decreases the production of pro-tumorigenic cytokines in co-culture. A) NISCH expression (green) in CAFs isolated from two patient tissues, phalloidin red, scale bar 20 μm; B) Percent viability of CAFs after 72 hours of treatment with increasing concentrations of rilmenidine, determined by MTT assay, presented as log(inhibitor) vs. normalized response; C) Immunofluorescence staining of CAF markers α-SMA (red), FAP (red), collagen I (green) and fibronectin (green) after 72 h of rilmenidine treatment in CAF1 (left panel) and CAF2 (right panel) cells, scale bar 50 μm; D) Expression of α-SMA, FAP and fibronectin in CAF1 and CAF2 cell lysates, after 72 h of rilmenidine treatment; E) Expression of α-SMA, FAP and fibronectin in CAF1 and CAF2 from co-cultures with either PANC-1 or MIA PaCa-2 cells, untreated or treated with 100 μM rilmenidine; F) mRNA expression of *ACTA2* (α-SMA), *FAP* and *FN1* in CAF2 cells alone or from co-cultures with PANC-1 and MIA PaCa-2 cells after rilmenidine treatment, normalized to untreated; G) Quantification of the integrated density of signal from the dot blot analysis of conditioned media from PANC-1 - CAF1 and PANC-1 - CAF2 co-cultures treated with 100 μM rilmenidine; H) Quantification of common cytokine profile changes from dot blot analysis of conditioned media from PANC-1 - CAF1 and PANC-1-CAF2 co-cultures treated with 100 μM rilmenidine. Mean ± SD, ****p < 0.0001, one-way ANOVA (Dunnett’s multiple comparisons test), n = 2; I) mRNA expression of select cytokines in CAF1 and CAF2 cells from co-cultures with PANC-1 cells treated with rilmenidine; J) mRNA expression of select cytokines in PANC-1 cells from co-cultures with CAF1 or CAF2 cells, compared to and normalized to untreated control. The graphs show mean ± SE, n = 3.

CAFs make up a barrier for chemotherapy and targeted therapy and can produce signaling molecules that can facilitate tumor progression and metastasis (67). The tumor microenvironment communicates with cancer cells via a variety of secreted growth factors and cytokines. We examined the cytokine profile of CAF-cancer cell co-cultures upon treatment with 100 μM rilmenidine by dot blot. While the cytokine profiles of the conditioned medium of CAF1 (**Fig. 6G left panel**) and CAF2 cells (**Fig. 6G right panel**) co-cultured with PANC- 1 cancer cell line differed, we observed common changes in a decrease of cytokines CCL2, IL-6 and IL-8 in the 100 μM rilmenidine treated group (**Fig. 6H**). ELISA assays on CAF2-cancer cell co-culture media confirmed decreased IL-6 and IL-8 secretion in co-culture media upon rilmenidine treatment (**Supplementary Fig. S5A and B**). PAI-1 levels were much higher in CAF-PANC-1 co-cultures compared to CAFs alone, but rilmenidine treatment did not alter its expression (**Supplementary Fig. S5C**).

Next, we examined the mRNA expression of cytokines and components of the fibrotic signature in CAFs and PANC-1 cells from co-cultures. We found a decrease in *IL6* and *PAI-1* mRNA levels in both CAF1 and CAF2 cells from co-cultures after 72 h of treatment with 100 μM rilmenidine (**Fig. 6I**), and a decrease in *IL8* mRNA level in the CAF2 cell line. *IL8* had a trend of decrease only in PANC-1 cells co-cultured with CAF2, while the PANC-1 cells from the co-culture with CAF1 showed a significant decrease in mRNA levels for *THSP* (**Fig. 6J**). Taken together, our results imply that although there was variability in response between CAFs from different patients, rilmenidine has an effect on the phenotype of CAFs from the TME, and that it has the potential to reduce communication between CAFs and cancer cells, which is the crucial process in PDAC progression.

## DISCUSSION

Although the five-year survival rate of patients with PDAC has surpassed 10% in the last decade, the increase in survival is still modest, and the incidence of the disease is on the rise (68). PDAC is fatal once it has spread beyond the primary site. With limited treatment options, the research focus is on discovering better therapy targets, especially for metastatic disease, and biomarkers that will enable earlier detection. The only potentially curative option for PDAC is surgery, preceded with neoadjuvant and/or followed by the adjuvant chemo/radiotherapy, but only 20% of patients are eligible for this type of treatment (69). Notwithstanding, in 75% of operated patients metastatic relapse occurs in less than 5 years (69,70). Mouse models of PDAC revealed that epithelial-mesenchymal transition in pancreatic cells occurs early, at the level of precancerous PanIN lesions, which leads to stroma and bloodstream invasion and potential seeding to distant organs (10). Analysis of clonal populations that give rise to distant metastases predicted that five years are required for the initial non-metastatic founder cell to acquire the metastatic ability (11). In PDAC patients who have undergone surgical resection, recurrence occurs most often in 12-24 months (71,72). This creates a window of opportunity to deliver potential antimetastatic therapy. Metastatic spread is a complex cascade of events that includes cell detachment, migration and local tissue invasion, intravasation into the vasculature or lymphatics, survival in the circulation, extravasation, attachment and adaptation to the new microenvironment, growth of micrometastases, and finally establishment of macrometastases (73). Targeting any of these processes may help limit PDAC progression.

NISCH has been identified as a tumor suppressor in breast and ovarian tumors, where the decrease of its expression is associated with disease progression (31,74). In breast cancer, the level of expression is also associated with cell invasiveness *in vitro*, where moderately invasive cells (MCF-7 and T47D) had a higher NISCH expression than the more invasive (MDA-MB-231) cells (74). NISCH inhibits cell migration, through binding to the integrin α5 and interaction with proteins that modulate cell-cell and cell-ECM adhesion and reorganization of the actin cytoskeleton (24,62). As control of adhesion and migration are prerequisites for cancer cell invasion and spread (75), NISCH is a potentially attractive antimetastatic drug target. There are three FDA- and European Medicines Agency (EMA)-approved antihypertensive drugs that bind to IRAS-I1 receptors: rilmenidine, moxonidine, and clonidine (34). Given that NISCH is an IRAS-1 protein (76), we investigated whether these drugs could be repurposed in the PDAC setting.

We have previously examined NISCH expression in datasets from the public repositories in various tumor types and reported that it was expressed to a greater or lesser extent in all the examined tumor tissues, most often decreased compared to the healthy counterparts (33). In this study, we show that NISCH is expressed in pancreatic cancer, but that there is no significant difference in expression between the tumor and the healthy pancreatic tissue. However, based on the expression level in tumors, *NISCH* was a positive prognostic marker for both the progression free and overall survival of PDAC patients. We found no difference between the sexes in *NISCH* expression or its prognostic value. Notably, we found that NISCH was expressed in both epithelial and stromal compartments of PDAC tissue, and in cancer cells and CAFs *in vitro*. As stroma plays a significant role in PDAC tumor progression and response to therapy (77–79), NISCH expression in both compartments implies that systemically administered NISCH agonists may have diverse effects.

We first examined the effect of NISCH agonists on PDAC cancer cell fitness *in vitro,* and found that rilmenidine was the most potent drug, which was in agreement with rilmenidine having the highest selectivity for binding to NISCH (80). Although rilmenidine ultimately induced cancer cell apoptosis, concentrations needed to induce this effect were much higher for PDAC cell lines than previously reported by our group and others, for melanoma cells (81) and K562 leukemia cells (82). These concentrations are also not achievable in patients (83,84). However, *in vitro* studies generally use supra-physiological concentrations of drugs that cannot be administered to patients. For example, in studies of metformin effects on cancer cells, it has been shown that much higher doses are needed in *in vitro* assays (85–91) than *in vivo* studies and in patients (92–94) to reach the comparable effects. In our study, there were differences in sensitivity to rilmenidine among the tested cell lines that were irrespective of their doubling time or the amount of NISCH expressed, modeling the heterogeneity. Transcriptome analysis of the most sensitive cell line MIA PaCa-2 showed enrichment of pathways associated with cell adhesion and migration after the treatment with rilmenidine, as expected from the NISCH function.

We next confirmed in *in vitro* assays that rilmenidine treatment delays and at a high dose prevents cancer cell attachment on various ECM substrates, and that it inhibits cell migration. This is in agreement with the findings that NISCH regulates focal adhesion and invadopodia formation (62). However, in the invasion transwell assay rilmenidine only slightly impaired invasion irrespective of the matrix coating, apart from the BxPC-3 cell line where inhibition was significant. Mesenchymal (dependent on adhesion and MMP activity) and amoeboid (independent of MMP activity) modes of movement through matrices are dictated by both cell intrinsic characteristics and the microenvironment through which they invade (95), and cancer cells can move through matrices even without any ECM adhesion (96). PDAC cells show phenotypic plasticity when it comes to the type of migration and can migrate both dependently and independently of MMP secretion (64), therefore effects of rilmenidine may be dispensable in quasimesenchymal cell lines PANC-1 and MIA PaCa-2, when they move through matrices without ECM adhesion. Examining the gelatinolytic activity we found that BxPC-3 cells expressed higher amounts of collagenases than the two other lines, and that rilmenidine slightly decreased their activity. Ultimately, to ascertain the anti-metastatic potential of rilmenidine, we examined its effect on metastasis formation in the zebrafish model. Tg(*fli1*:EGFP) zebrafish, in which the promoter for the endothelial marker *fli1* drives the expression of enhanced green fluorescent protein (EGFP) in blood vessels, enables easy visualization and quantification of metastatic cancer cell seeding (97,98). In the PANC-1 tumor xenograft zebrafish model, all the embryos developed metastasis, with an average number of 6. Treatment with 50 μM rilmenidine significantly reduced the number of metastases per embryo, and at 100 μM more than 70% of embryos had no detectable metastases. These results imply that rilmenidine is effective in decreasing the metastatic spread.

As the main feature of PDAC is the prominent TME, it is imperative to examine the effects of the treatment on as many cell types as possible that are present in the tumor. Cancer cells and CAFs are in dynamic crosstalk through direct cell-to-cell contact and secretion of growth factors, cytokines, and extracellular vesicles. This bi-directional communication can have both tumor-promoting and tumor-suppressive roles in PDAC development (99). CAFs can promote migration and invasion of cancer cells, as well as the drug resistance (67), and can create a pro-inflammatory microenvironment (17). Several distinct CAF phenotypes have been identified, out of which the three main types – myofibroblasts (myCAFs), inflammatory CAFs (iCAFs), and antigen-presenting CAFs (apCAFs) – are the most well described. MyCAFs are mostly located adjacent to neoplastic cells and are characterized by high expression of αSMA and lower expression of FAP, iCAFs express FAP and secrete cytokines such as IL-6 that support the growth of cancer cells (17), while apCAFs express the MHC II complex and CD74 and can present antigens to T cells *in vitro* (100). CAF phenotypes are not fixed and subtypes can transition between subsets (101). We were able to isolate mixed populations of CAFs from three patients and examine the effects of rilmenidine on both monocultures and cancer cell-CAF co-cultures. NISCH was expressed in CAF cells, but even high doses of rilmenidine of up to 600 μM did not affect their viability. However, 100 μM rilmenidine treatment decreased both αSMA and FAP markers, and to an extent deposition of the ECM. This held true in CAFs co-cultured through a transwell system with pancreatic cancer cells. These findings are of importance, as they imply that rilmenidine may contribute to the normalization of the tumor stroma. We further examined the effect of rilmenidine treatment on cytokine production in co-culture. Rilmenidine treatment significantly decreased the production of IL-6, IL-8, and CCL-2 in co-cultures. These results are promising, as IL-6 is an important player in PDAC progression (102–104). IL-6 secreted by CAFs stimulates the activation of STAT3 signaling in tumor cells (105) and promotes invasion (106). IL-6/STAT3 signaling also contributes to tissue fibrosis (107), and inhibition of IL-6 signaling showed promising results in the inhibition of pancreatic tumor progression (108). We have also observed a decrease in PAI-1 production in co-cultures, as well as a decrease in transcription of *THSP*, another fibrosis marker, in both CAFs and cancer cells. Taken together, these results imply that rilmenidine treatment may have positive effects on tumor stroma, by attenuating CAF pro-tumorigenic phenotype, a decrease of fibrotic signature and disruption of cancer cell-CAF communication.

In conclusion, our study lays the ground for repurposing the antihypertensive rilmenidine in the treatment of PDAC, as it targets at least two aspects of PDAC progression – cancer cell dissemination and cancer cell-stroma communication. However, further examination of rilmenidine effects in *in vivo* PDAC models is needed to reassess the issue of the dosing and examine the effects on other important aspects of the disease, such as the effect on tumor vasculature and immune infiltrate. While repurposing rilmenidine would provide faster translation to the clinic, the design and development of novel NISCH agonists may also be of interest, as we show that NISCH is expressed in both tumor and the stromal compartments and its agonization may have a dual effect.

## Clinical perspectives

- Nischarin (NISCH) is a scaffolding protein with tumor suppressive role; however, its role and the effects of its activation have not been investigated in pancreatic ductal adenocarcinoma (PDAC).
- NISCH was expressed in both the epithelial and stromal compartments of PDAC. NISCH agonist rilmenidine had an effect on the migration and invasion of tumor cells, and the secretory functions of both cancer cells and cancer associated fibroblasts (CAFs). Additionally, rilmenidine successfully reduced the metastatic spread of PDAC cells in an *in vivo* xenograft zebrafish model.
- Our results suggest that rilmenidine is a good candidate for drug repurposing and that NISCH is a promising target for the development of new, more selective agents to treat PDAC patients.

## Acknowledgements

All authors have read the journal’s policy on disclosure of potential conflicts of interest and all authors have no financial or personal relationship with organizations that could potentially be perceived as influencing the described research.

## CRediT authorship contribution statement

KZ performed experiments, analysed data, and wrote the original manuscript draft. MP performed experiments, analysed data and reviewed and edited the manuscript. MO curated-analysed data and reviewed and edited the manuscript. DG managed patients and provided PDAC patient tissues. AP conducted zebrafish experiments, analysed data and reviewed and edited the manuscript. TSR performed experiments and reviewed and edited the manuscript. JG conceptualised the study, designed experiments, provided project supervision, funding acquisition, and reviewed and edited the manuscript. All authors read and approved the final manuscript.

