## Supplemental Material and Figures for "Nischarin agonist rilmenidine shows antimetastatic potential in pancreatic ductal adenocarcinoma"

**Supplementary Table S1. List of assays performed on CAF cell lines.**

| *Experiment* | *CAF1* | *CAF2* | *CAF3* |
|---|---|---|---|
| *ICC NISCH* | + | + | - |
| *MTT assay* | + | + | - |
| *ICC fibroblast markers* | + | + | + |
| *Monoculture WB* | + | + | + |
| *Co-culture WB* | - | + | + |
| *Co-culture qPCR* | + | + | - |
| *Dot blot* | + | + | - |

**Supplementary Table S2. Lethal and teratogenic effects observed in zebrafish (*Danio rerio*) embryos at different hours post fertilization (hpf).**

| *Category* | *Toxicological parameters* | *Exposure time (hpf)* | | | | |
|---|---|---|---|---|---|---|
| | | **24** | **48** | **72** | **96** | **120** |
| *Lethal effect* | Coagulated eggs[a] | ● | ● | ● | ● | ● |
| | Lack of the heart beating | ● | ● | ● | ● | ● |
| | Non-detachment of the tail | ● | ● | ● | ● | ● |
| | Lack of somite formation | ● | ● | ● | ● | ● |
| *Teratogenic effect* | Malformation of head | ● | ● | ● | ● | ● |
| | Malformation of eyes[b] | ● | ● | ● | ● | ● |
| | Malformation of sacculi/otoliths[c] | ● | ● | ● | ● | ● |
| | Malformation of chorda | ● | ● | ● | ● | ● |
| | Malformation of tail[d] | ● | ● | ● | ● | ● |
| | Scoliosis | ● | ● | ● | ● | ● |
| | Yolk edema | ● | ● | ● | ● | ● |
| | Yolk deformation | ● | ● | ● | ● | ● |
| | Growth retardation[e] | | ● | ● | ● | ● |
| | Hatching | | | ● | ● | ● |
| | Swimmbladder development | | | | | ● |
| *Hepatotoxicity* | Yolk absorption | | | ● | ● | ● |
| | Liver darkening | | | ● | ● | ● |
| *Cardiotoxicity* | Pericardial edema | | ● | ● | ● | ● |
| | Heart morphology | | | ● | ● | ● |
| | Heart beating rate (beat/min) | | | | ● | ● |

[a]No clear organs structure is recognized

[b]Malformation of eyes was recorded for the retardation in eye development and abnormality in shape and size.

[c]Presence of none, one or more than two otoliths per sacculus, as well as reduction and enlargement of otic vesicles

[d]Tail malformation was recorded when the tail was bent, twisted or shorter than to control embryos as assessed by optical comparison

[e]Growth retardation was recorded by comparing with the control embryos in a body length (after hatching)

**Supplementary Table S3. Effect of nischarin agonists on viability of the pancreatic cancer cell line panel.**

| ***Cell line*** | ***$IC_{50}$ (μM)*** | | |
|---|---|---|---|
| | **Rilmenidine** | **Clonidine** | **Moxonidine** |
| ***PANC-1*** | 306.8 | >1000 | >1000 |
| ***MIA PaCa-2*** | 169.2 | >1000 | 809.4 |
| ***BxPC-3*** | 367.7 | >1000 | 320.1 |
| ***Capan-2*** | >1000 | >1000 | >1000 |
| ***HPAF-II*** | 408.9 | >1000 | ND |
| ***CFPAC-1*** | 579 | >1000 | ND |
| ***SW1990*** | 458.9 | >1000 | ND |
| ***AsPC-1*** | 968.9 | ND | ND |

***ND = not determined**

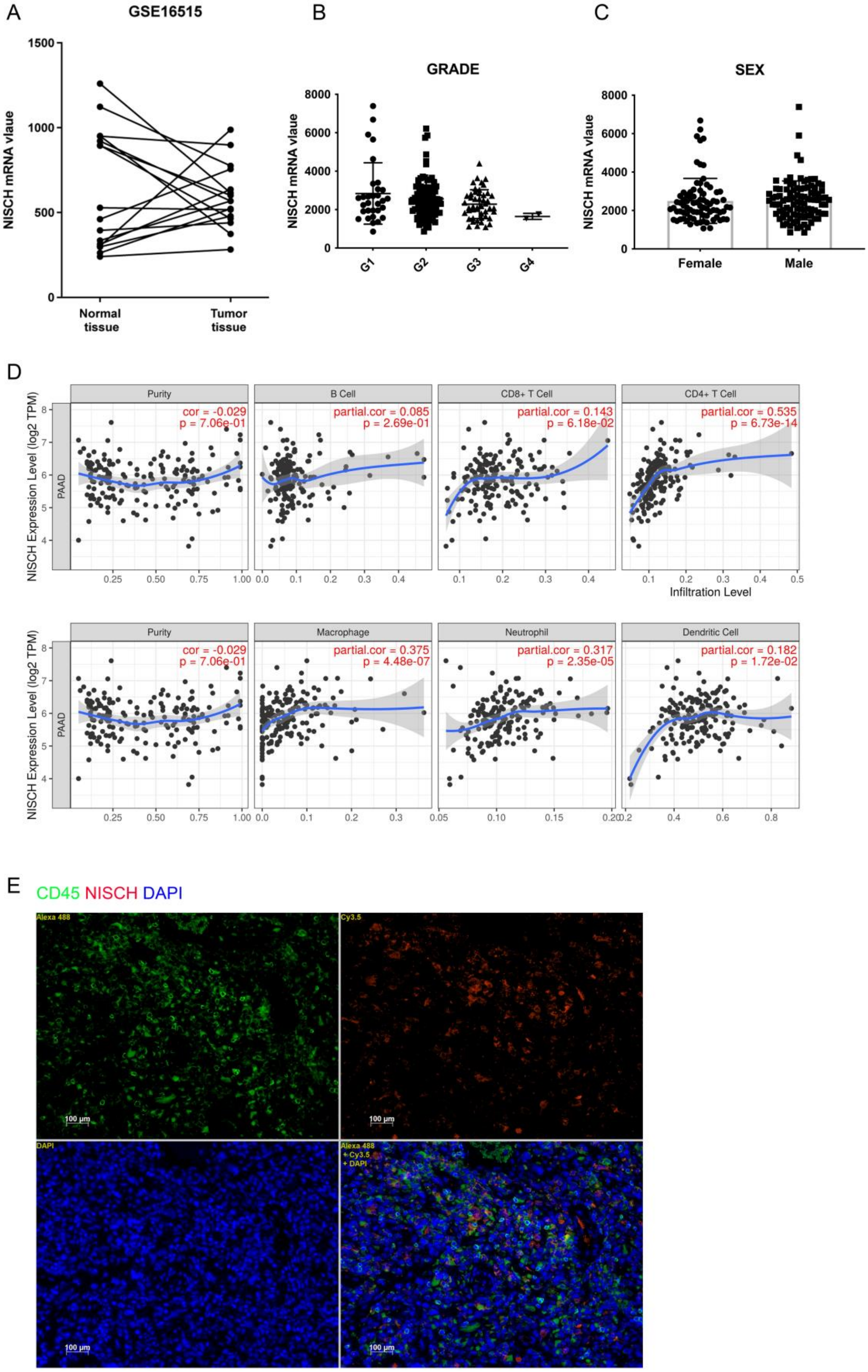


**Supplementary Figure S1.** A) *NISCH* mRNA expression in the paired tumor and adjacent PDAC tissue samples in the GSE16515 cohort; B) *NISCH* mRNA expression by grade in the

TCGA PAAD cohort C) *NISCH* mRNA expression by patient sex in the TCGA PAAD cohort; D) Correlation of *NISCH* expression with immune infiltration level in PDAC, shown are purity-corrected partial Spearman's rho value and statistical significance (Li et al. 2017) E) Expression of NISCH (red) and CD45 (green) in PDAC liver metastasis sample from the NBP2-78128 microarray, nuclei DAPI blue, scale bar 100 µm .**Supplementary Figure S2.** A) Quantification of PANC-1, B) MIA PaCa-2 and C) BxPC-3 cell spreading over time, in presence of increasing concentrations of rilmenidine on tissue culture plastic, collagen I and fibronectin coated surfaces. Mean ± SEM. Two-way ANOVA (Sidak's multiple comparisons test), $n = 3$; *$p < 0.05$, **$p < 0.01$, ***$p < 0.001$, ****$p < 0.0001$. D) Representative images of PANC-1, E) MIA PaCa-2 and F) BxPC-3 cell spreading (left panel) and wound closure (right panel) for after 24 hours of rilmenidine treatment. Magnification: 10x.

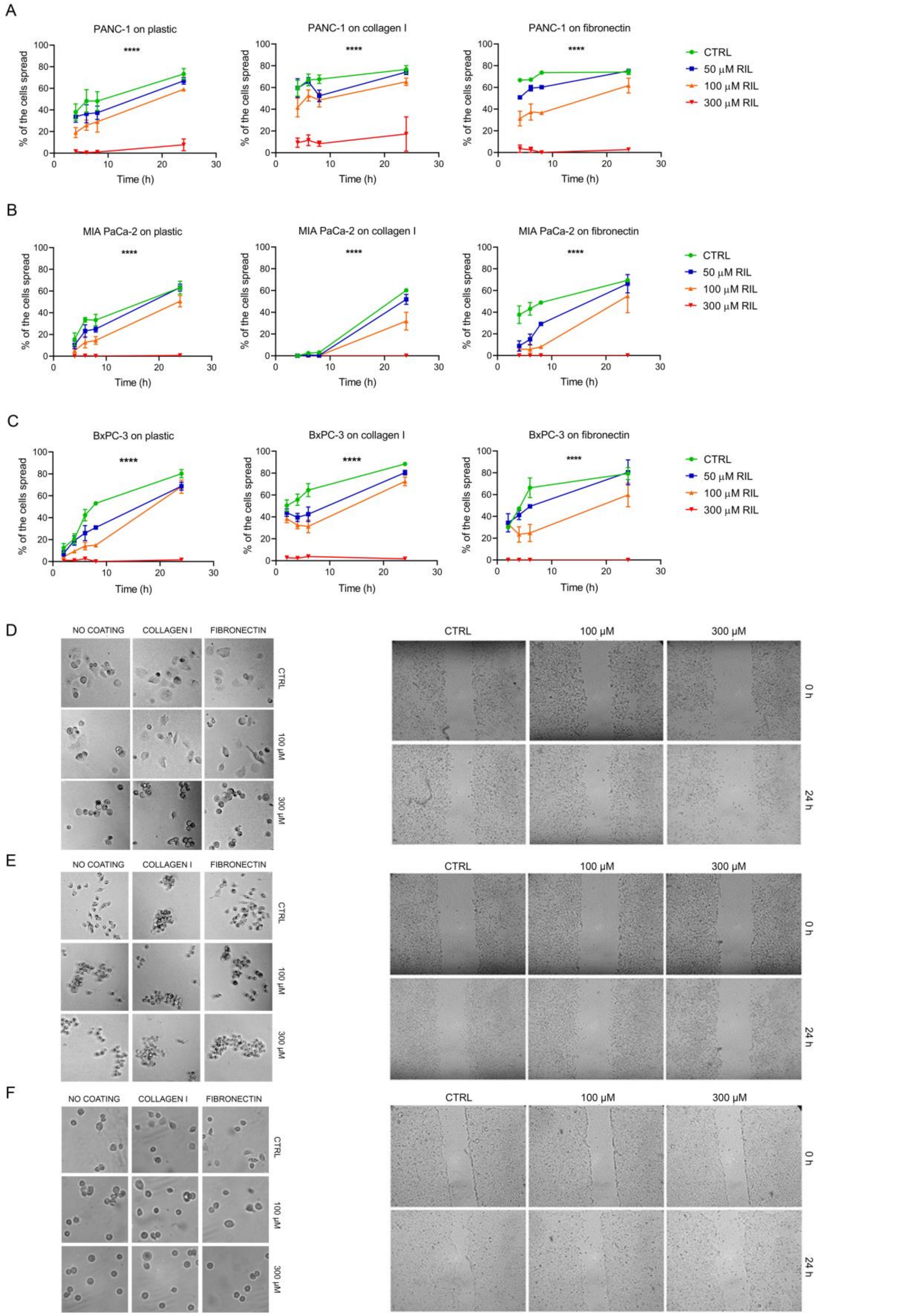


**Supplementary Figure S2.** A) Quantification of PANC-1, B) MIA PaCa-2 and C) BxPC-3 cell spreading over time, in presence of increasing concentrations of rilmenidine on tissue culture plastic, collagen I and fibronectin coated surfaces. Mean ± SEM. Two-way ANOVA (Sidak's multiple comparisons test), $n = 3$; *$p < 0.05$, **$p < 0.01$, ***$p < 0.001$, ****$p < 0.0001$. D) Representative images of PANC-1, E) MIA PaCa-2 and F) BxPC-3 cell spreading (left panel) and wound closure (right panel) for after 24 hours of rilmenidine treatment. Magnification: 10x.

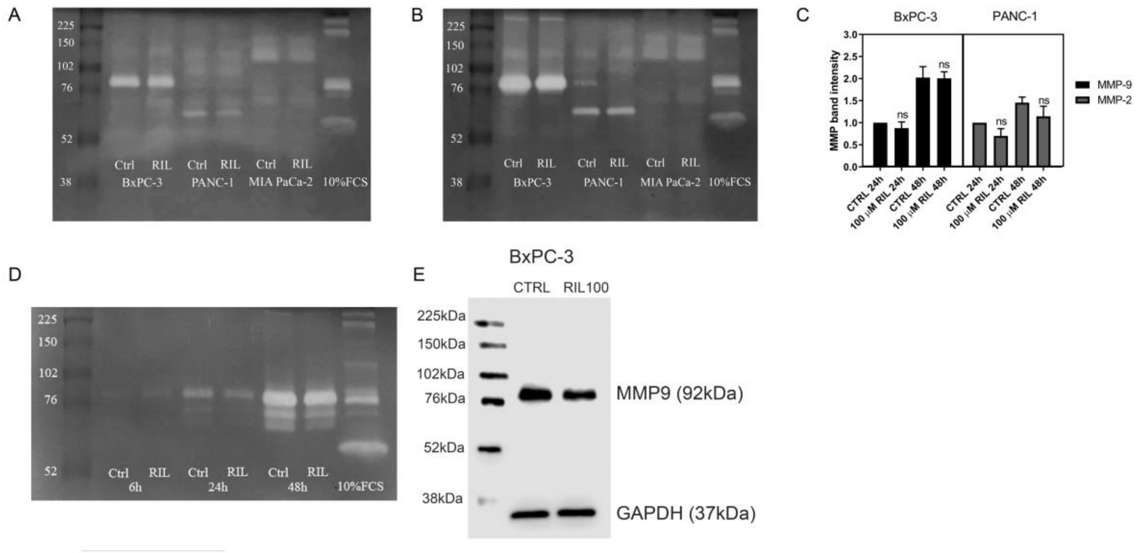


**Supplementary Figure S3. Rilmenidine effects on the gelatinolytic activity of MMP-2 and MMP-9 in PDAC cells *in vitro*.** A) Representative gelatin zymograms of 24 h or B) 48 h conditioned medium of PANC-1, MIA PaCa-2, and BxPC-3 cells treated with 100 µM rilmenidine C) Quantification of the intensities of cleared bands analyzed with ImageJ and normalized to untreated. Mean ± SD, Two-way ANOVA (Sidak's multiple comparisons test), n = 3; (compared to the untreated control). D) Gelatin zymography of BxPC-3 cell lysates 6 h, 24 h, and 48 h after the treatment with 100 µM rilmenidine. (D) MMP-9 expression in BxPC-3 cell lysates after 48 h of rilmenidine treatment.

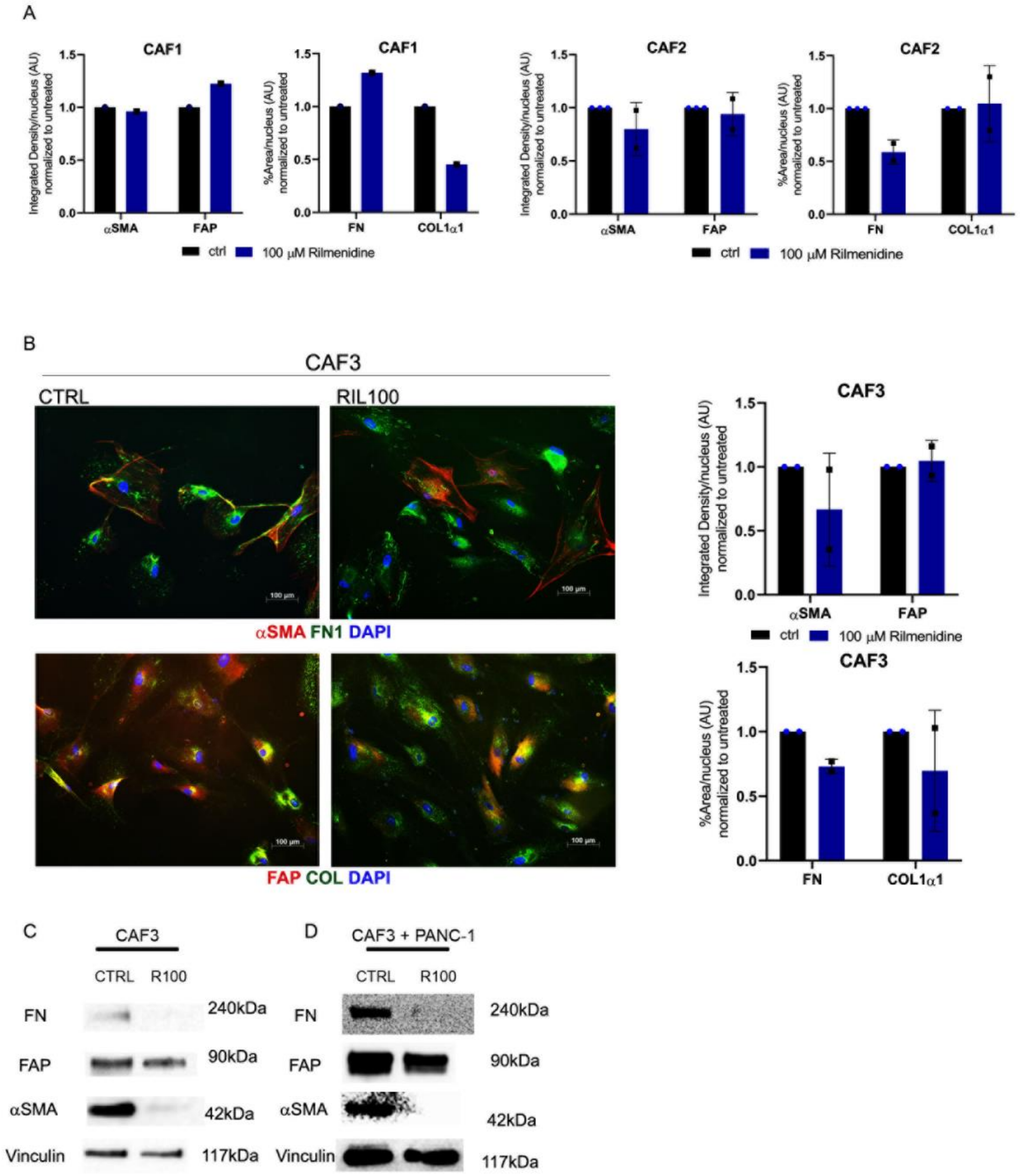


**Supplementary Figure 4.** A) Quantification of the immunofluorescence staining with Image J software of CAF markers α-SMA, FAP, collagen I and fibronectin after 72 h of rilmenidine treatment in CAF1 (left panel) and CAF2 (right panel) presented as integrated density of red or green signal per nucleus. B) Immunofluorescence staining of CAF protein markers α-SMA (red), FAP (red), collagen I (green) and fibronectin (green) after 72 h of rilmenidine treatment in CAF3 cells, scale bar 100 μm (left panel) and quantification of integrated density of staining per nucleus, normalized to untreated controls (right panel). C) Expression of α-SMA, FAP and fibronectin in CAF3 cells untreated or treated with 100 μM rilmenidine. D) Expression of α-SMA, FAP and fibronectin in CAF3 cells from co-cultures with PANC-1 cells, untreated or treated with 100 μM rilmenidine.

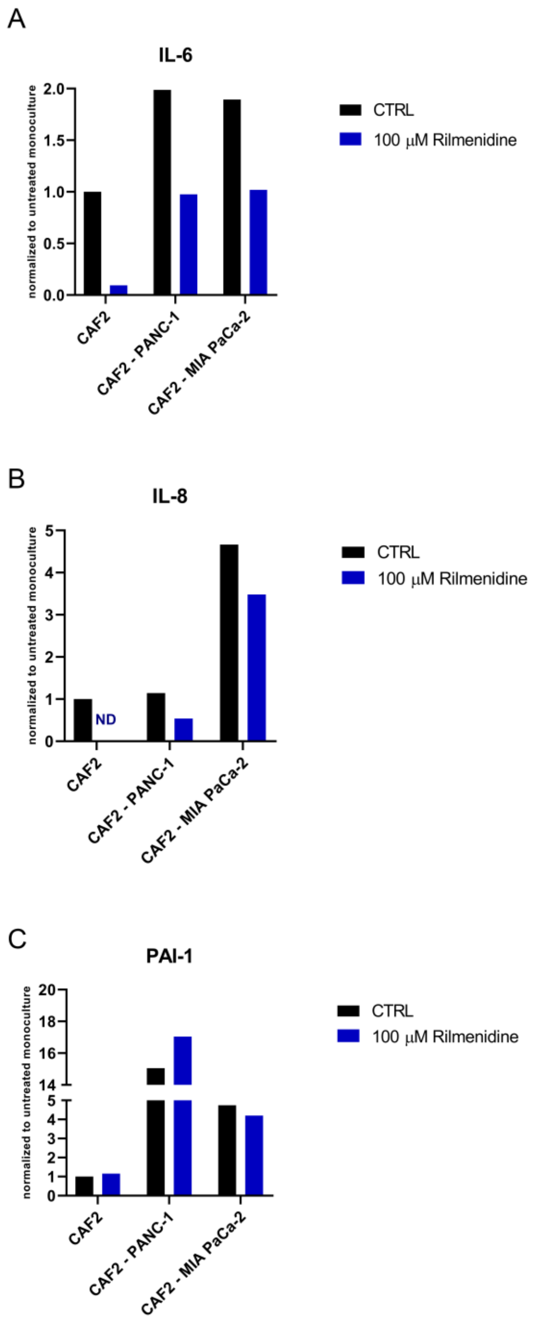


**Supplementary Figure S5. Rilmenidine impacts the level of cytokine production in co-cultures and patient tissues.** Levels of A) interleukin 6, B) interleukin 8 and C) plasminogen activator inhibitor 1 in culture media of CAFs and pancreatic cancer cells co-cultures after treatment with 100 μM rilmenidine.
